# Pathogenic variants of the mitochondrial copper chaperones SCO1 and SCO2 reshape their respective interactomes and disrupt cellular phospholipid metabolism

**DOI:** 10.64898/2026.09.23.753947

**Authors:** Sampurna Ghosh, Zakery N. Baker, Rachel M. Guerra, Hana Antonicka, Abhinav B. Swaminathan, Vishal M. Gohil, David J. Pagliarini, Stanley A. Moore, Scot C. Leary

**Author notes:** Corresponding authors: Scot C. Leary, 3069664349, Stanley A. Moore, 3069664381.

## Abstract

SCO1 and SCO2 are required for copper delivery to COX2, a copper-containing subunit of cytochrome *c* oxidase (COX), yet how mutations in these genes cause distinct, tissue-specific forms of human disease remains poorly understood. To gain further insight into the molecular underpinnings of this clinical heterogeneity, we used BioID to map the interactomes of four pathogenic SCO variants (SCO1 G132S, SCO1 P174L, SCO1 M294V and SCO2 E140K) and the wild-type proteins. While this approach identified many proteins common to both wild-type neighbourhoods, several potential interacting partners unique to each SCO protein were also observed that were consistent with their known roles in COX assembly. Follow-up analyses revealed that SCO1 interacts with COX16 and that this interaction is stabilized within the membrane by a coiled-coil helix-helix interface, with the soluble C-terminal region of COX16 physically bridging SCO1 and COX2 within a ternary complex to facilitate copper delivery. We further observed that COX16 abundance is relatively low in the brain and its association with SCO1 is most severely impaired by the M294V substitution associated with a fatal encephalopathy. Intriguingly, our BioID analyses also detected significant enrichment in each SCO neighbourhood for biosynthetic enzymes and lipases critical to phospholipid metabolism and found that the affinity for these candidate interactors was uniquely perturbed by various pathogenic variants of SCO1 and SCO2. Collectively, our data emphasize the potential of proximity labelling to further define the molecular roles of disease-causing variants that perturb mitochondrial function and suggest that SCO proteins impinge upon phospholipid metabolism.

## Introduction

Mitochondria are comprised of approximately 1,100 proteins of dual genetic origin (1–3) and make a myriad of fundamental contributions to cellular homeostasis (4, 5). Mutations in protein products from either genome impair organelle function and result in human disease (6–8), with mitochondrial disorders as a collective representing the largest group of inborn errors of metabolism (9–11). The central role of mitochondria in cellular physiology is underscored by the broad spectrum of clinical phenotypes associated with mitochondrial disorders, which can affect virtually any organ system of the body and complicates diagnosis, obscures genotype–phenotype relationships and presents a major challenge for the development of effective therapeutic interventions (6, 12).

Among mitochondrial diseases reported to date, isolated cytochrome *c* oxidase (COX) deficiencies are a rare but significant cause of respiratory chain defects (13, 14). COX is comprised of fourteen structural subunits and is assembled in a linear-modular manner (15, 16), with the nuclear-encoded subunits surrounding the mitochondrially-encoded enzymatic core which contains modified heme and copper prosthetic groups essential for catalytic competence (17). Mutations that impair the temporally coordinated maturation or stoichiometric integration of mitochondrially- and nuclear-encoded subunits into the nascent holoenzyme complex often lead to severe, tissue-specific forms of disease. Deleterious mutations in mitochondrially-encoded COX subunits typically manifest as encephalomyopathy, myopathy, exercise intolerance, optic atrophy, retinopathy and hearing loss in late childhood or early adulthood owing to the multi-copy nature of the mitochondrial genome (14, 18–20). In contrast, the autosomal inheritance of pathogenic mutations in nuclear genes encoding structural COX subunits or ancillary factors that facilitate holoenzyme assembly result in severe, early onset forms of tissue-specific disease that are ultimately fatal (13).

A considerable number of COX assembly factors are essential for the maturation of the COX2 subassembly module and, more specifically, the metalation of its Cu_A_ site (21, 22). The biogenesis of this mixed valence, binuclear prosthetic group located within a solvent-exposed thioredoxin fold in the mitochondrial intermembrane space (IMS) relies minimally upon SCO1, SCO2, COX16, COA6 and COA7 (23–28), with COX17 acting as the primary upstream copper donor (29, 30). SCO1 and SCO2 are evolutionarily conserved paralogs that were originally identified in yeast as high copy suppressors of a *COX17* point mutant (31). Both Cu(I) and Cu(II) binding are essential for SCO protein function (32), with SCO2 acting as a thiol-disulfide oxidoreductase to help prime two copper-coordinating cysteines of COX2 for subsequent metalation by SCO1 (33, 34). With few exceptions (35, 36), pathogenic mutations in either *SCO* gene result in a severe, combined COX and copper deficiency associated with early onset forms of fatal, tissue-specific disease (37, 38).

*SCO2* mutations have been identified in roughly 100 pedigrees with almost all patients carrying at least one E140K substitution and presenting primarily with a fatal, neonatal encephalocardiomyopathy (39–42). In contrast, *SCO1* mutations are very rare and thus far only four pedigrees with early onset, fatal forms of clinically heterogeneous disease have been reported. The first *SCO1* patient was a compound heterozygote carrying a nonsense mutation on one allele and a P174L substitution on the other allele who presented with neonatal liver failure (43). The second *SCO1* patient was homozygous for a G132S substitution and died at 6 months of age due to hypertrophic cardiomyopathy (38). The third *SCO1* patient was a compound heterozygote harboring a nonsense mutation on one allele and an M294V substitution on the other allele who died at 5 months of age from an isolated encephalopathy (44). Finally, the fourth *SCO1* patient was homozygous for a G106del mutation and died at 1 month of age from chronic lactic acidosis in the absence of a detectable hepatopathy, encephalopathy or cardiomyopathy (45).Why mutations in these ubiquitously expressed paralogs operating within the same biochemical pathway result in strikingly different, tissue-specific clinical phenotypes is a phenomenon that remains poorly understood.

Here, we hypothesize that SCO protein function impinges upon additional, non-canonical pathways and that pathogenic variants of SCO1 and SCO2 perturb their respective interactomes such that it is particularly detrimental to tissue function when interacting partners are present at low abundance. To address this possibility, we used BioID to systematically map the interactomes of wild-type SCOs and selected pathogenic variants. We further assessed the relative abundance of potential interacting partners in heart, liver and brain by mining existing proteomics data. Our efforts led to the identification of a small subset of proteins that conform to our hypothesis, most notably COX16 whose interaction with SCO1 is essential for its stability and severely disrupted by the SCO1 M294V substitution associated with a fatal encephalopathy. Our data also uncovered an unexpected enrichment in SCO interacting partners critical to phospholipid metabolism. Collectively, our findings emphasize the power of using proximity labelling studies to further define the molecular roles of disease-causing variants that perturb mitochondrial function.

## Materials and Methods

### Cell lines

Flp-In T-REx 293 cells (Invitrogen) were maintained in high-glucose DMEM (Gibco) supplemented with 10% fetal bovine serum (FBS) and 2mM glutaMAX (Gibco) in an atmosphere of 5% CO_2_ at 37°C. At 70% confluency, cells were co-transfected with pDEST-pcDNA5-BirA*-FLAG-C-terminal bait (46) or spatial control constructs and pOG44 vector using Lipofectamine 2000, according to the manufacturer’s instructions. Stable integrants were selected in media containing 200 U/mL hygromycin B, and single cell clones were isolated by serial dilution. Four to six clones per construct were then characterized to ensure comparable bait protein expression, subcellular localization and biotinylation activity.

Primary fibroblasts from control individuals and patients carrying pathogenic variants in *SCO1*, *SCO2*, *COA6* or *COX16* were immortalized as previously described (47, 48). Fibroblasts were retrovirally transduced with constructs of interest (25, 28) using the Phoenix amphotropic packaging cell line (a kind gift of Garry P. Nolan) and selected with hygromycin B or puromycin. A *COX16* cDNA was amplified from reverse transcribed total RNA with 5’-ACAAAAAAGCAGGCTATGTTTGCACCCGCGGTGATG-3’ and 5’-ACAAGAAAGCTGGGTTCAAGTTGTCTTAGTCTTAAGGCTTTCTGG-3’, cloned into pDNOR221 (Invitrogen), Sanger sequenced and transferred to a retroviral overexpression vector. Floxed and *Sco1* null mouse embryonic fibroblasts (MEFs) (49) were transduced with retroviral vectors containing a wild-type *Sco1* cDNA or the murine equivalents of the pathogenic human variants (50), and selected with hygromycin B. All cell lines were routinely tested and confirmed to be free of mycoplasma contamination.

### BioID construct design and cloning

Wild-type and mutant human *SCO1* and *SCO2* cDNA sequences were amplified from existing retroviral expression vectors (25, 44) and cloned into pDONR221. Fidelity was verified by Sanger sequencing, and a hydrophilic linker sequence (DPKESGSVSDSR) (51) was introduced between each cDNA and the BirA* portion of the fusion protein by PCR prior to Gateway transfer into the pDEST-pcDNA5-BirA*-FLAG-C-term destination vector. The mitochondrial targeting sequences of OPA1 or AIFM1 were also PCR amplified and cloned into the pDEST-pcDNA5-BirA*-FLAG-C-term vector and used as spatial controls to account for background biotinylation within the mitochondrial intermembrane space (IMS) (52). BirA-FLAG-GFP* was included to detect baseline biotinylation within the cytosol, while untransduced Flp-In T-REx cells were used to detect endogenously biotinylated proteins (52).

### BioID sample preparation

Protein expression was induced when cells were ∼70% confluent with 1μg/mL tetracycline and 50 μM biotin. Cells from duplicate 150 mm plates were then harvested 24 hours later in ice-cold PBS, pelleted by centrifugation at 1,000 *× g* for 5 min at 4 °C, and lysed in BioID lysis buffer (50 mM Tris [pH 7.5], 150 mM NaCl, 1% IGEPAL CA-630, 0.4% SDS, 1.5 mM MgCl_2_, 1 mM EGTA, 1X PIC) containing benzonase (Millipore). Lysates were freeze–thawed and placed on a rotor at 4°C for 30 mins to facilitate nucleic acid digestion, sonicated on ice (65% amplitude using three 10 second cycles with 2 seconds rest between cycles), and clarified by centrifugation at 17,000 *× g* for 20 min at 4°C. Protein concentrations were determined using the DC Protein Assay (BioRad), and a fraction was retained for validation via immunoblotting. Clarified lysates were then incubated overnight at 4°C with streptavidin-coated magnetic beads (Cytiva). Beads were washed sequentially with BioID wash buffer (50 mM HEPES [pH 8.0], 100 mM KCl, 10% glycerol, 2mM EDTA and 0.1% NP-40) and BioID lysis buffer, exchanged into 100 mM ammonium bicarbonate, and subjected to on-bead tryptic digestion followed by mass spectrometry.

### BioID mass spectrometry (MS), data processing and statistical analysis

The MS protocol for sample analysis was provided by Dr. Denis Faubert (Institut de Recherches Cliniques de Montréal). The protein-bead conjugates were digested overnight at 37°C by agitation with 0.5 μg Sequencing Grade Modified Trypsin (Promega). Supernatants were collected upon pelleting the beads, and the beads were washed two times with 100 μL of water to collect any residual peptide. All supernatants and associated washes were then pooled, reduced and alkylated. The reduction step was done with 9 mM DTT at 37°C for 30 min followed by cooling for 10 min at RT, while the alkylation step was done with 17 mM iodoacetamide at room temperature for 20 min in the dark. The supernatants were acidified with trifluoroacetic acid for desalting and residual detergents were removed by MCX (Waters Oasis MCX 96-well Elution Plate), following the manufacturer’s instructions. After elution in a 10% ammonium hydroxide/90% methanol (v/v) solution, samples were dried with a speed vac, reconstituted under agitation for 15 min in 15 µL of 5% formic acid and loaded onto a 75 μm i.d. × 150 mm Self-Pack C18 column installed in the Proxeon 1200 system (Thermo Scientific). The buffers used for chromatography were 0.2% formic acid (buffer A) and 85% acetonitrile/0.2% formic acid (buffer B). Peptides were eluted with a three-slope gradient at a flow rate of 250 nL/min. Solvent B increased initially from 4 to 20% over 45 min, then from 20 to 37% over 62 min and finally from 37 to 70% over 8 min.

The HPLC system was coupled to a Q Exactive mass spectrometer (Thermo Scientific) through a Nanospray Flex Ion Source. Nanospray and S-lens voltages were set to 1.3-1.8 kV and 50 V, respectively. Capillary temperature was set to 250°C. Full scan MS survey spectra (m/z 360-2000) in profile mode were acquired in the Orbitrap with a resolution of 70000, an AGC target set to 1^e6,^ and a maximum ion time set to 100 ms. The 15 most intense peptide ions were fragmented in the collision cell and MS/MS spectra were analyzed in the Orbitrap with a resolution of 17500, an AGC target set to 1^e5^, a maximum ion time set to 50 ms and the dynamical exclusion set to 8 seconds. Post run .mzXML files were generated from Thermo RAW files using the ProteoWizard converter (53), and implemented within ProHits (--filter “peakPicking true2”--filter “msLevel2”) as previously described (52). Significance Analysis of INTeractome (SAINT) analysis was used to calculate the probability of a true interactor via spectral counts and Bayesian statistics, and was performed using ProHits-viz version 6.0.4. A threshold set at a 1% Bayesian false discovery rate (BFDR) was used to select for high-confidence interactions as previously described (52).

### Prioritization of candidate interacting partners

The SAINT analysis generated candidate interactors for all SCO bait proteins, which were further refined using an in-house analytical pipeline (Supplementary Table 1). An initial, basic filter of BFDR ≤ 0.2 and SAINT score ≥ 0.73 was applied to align with parameters from a prior BioID study that included SCO1 as a bait (54). To account for differences in bait expression and the absence of load normalization before mass spectrometry, prey fold change values were divided by BirA* spectral counts as an internal measure of bait protein abundance. Fold change values of the filtered prey proteins were then compared to the IMS spatial controls AIFM1-BirA* and OPA1-BirA* to distinguish specific interactions from background. Fold change was calculated as the ratio of spectral counts in bait purifications to those in controls, with a small constant added to avoid division by zero. Only prey proteins with a fold enrichment ≥ 1 relative to the most enriched spatial control were retained. The remaining candidates were then cross-referenced with MitoCarta 3 (3) to identify mitochondrial proteins (Localization filter 1) and only those proteins localized to the OMM, IMS or IMM were further considered (Localization filter 2). For pathogenic variants, prey protein fold changes were normalized to the corresponding wild-type bait to derive a relative affinity score. Tissue-specific abundances (heart ventricle, heart atrial, liver, brain cerebellum, brain cortex) were obtained from the human proteome map (55), normalized using a z-transformation and visualized using a color-coded heatmap. For proteins lacking expression data in individual tissues, abundance was assigned as a minimum value before normalization. Proteins lacking expression data for all tissues were removed from the heatmap and included as a dot plot.

### Mitopathway Enrichment Analysis

Mitopathway enrichment analysis was performed on each SCO dataset. Mitopathway assignments for each protein were obtained from MitoCarta 3.0 (3). Fisher’s exact test was performed to calculate the enrichment and significance (p-value) for every term present in the dataset. Log_10_ FDR adjusted p-values were calculated using the Benjamini-Hochberg procedure to control for multiple hypothesis testing. Only pathways with an FDR under 20% were retained within the plot.

### Immunoblotting

Cultured fibroblasts and Flp-In T-REx cells were harvested and lysed in RIPA buffer to generate clarified extracts. Equal amounts of protein (10–30 μg) were resolved on appropriate precast polyacrylamide gels (Bio-Rad) and transferred to nitrocellulose membranes under semi-dry conditions. For the detection of biotinylated proteins, membranes were blocked in TBST (25 mM Tris [pH 7.4], 137 mM NaCl, 2.5 mM KCl and 0.05% Tween 20) containing 5% BSA for 4 hours at room temperature and incubated overnight with streptavidin–HRP (Invitrogen). Membranes were washed three times with PBS, incubated briefly in adult bovine serum blocking buffer (PBS containing 10% v/v adult bovine serum and 1% w/v Triton X-100) for 5 mins, washed again, and processed for imaging. All other membranes were blocked and probed with primary and secondary antibodies in TBST containing 5% BSA or 5% milk (Supplementary Table 2), with six TBST washes between steps. Proteins were visualized using enhanced chemiluminescence and imaged on a Bio-Rad ChemiDoc MP system.

### Immunofluorescence

Cells were seeded at 15% confluency on coverslips in 6-well plates and expression of the fusion protein was induced the following day with 1 μg/mL tetracycline. After 24 hours, cells were washed with PBS, fixed in 4% paraformaldehyde (Sigma) for 10 min, and permeabilized with 0.2% Triton X-100 for 5 min. Cells were then incubated with primary antibodies in PBS containing 3% BSA for 45–60 min at 37 °C, followed by Alexa-conjugated secondary antibodies under the same conditions (Supplementary Table 2). Coverslips were mounted with ProLong Diamond Antifade Mountant containing DAPI and imaged using a Zeiss LSM700 confocal microscope. Images were processed using ZEISS Zen Black software.

### In silico structural modeling and bioinformatics

All sequences were retrieved from NCBI and aligned with known homologs from mouse, beluga, zebrafish and yeast using Clustal Omega (56). The AlphaFold3 server (https://AlphaFoldserver.com) was used to model all structures and/or molecular complexes of interest (57, 58), both in the apo- and copper-loaded forms. In each case, the top ranked of five models is shown and recapitulation of their main features was confirmed by generating analogous models using murine and other vertebrate sequences (data not shown). For SCO1, COX16, COX2 and COX20, the extent of membrane spanning α-helices appeared to be relatively accurate and were consistently located within a spatial region likely to contain a membrane bilayer. In addition, we compared the locations of membrane spanning helices in each protein with the membrane-spanning region of COX2 in the structure of mature bovine COX (59). After selection of the best models, the complex structures were manually adjusted in Coot if necessary (side chain adjustments mostly and Cu-ligand residues) and then refined in PHENIX using standard protein parameters and known bond-lengths of Cu-ligand residues Cys, His and Met (60–62). Comparison of the atomic coordinates for various complexes was carried out using the SSM superpose function in Coot. More detailed comparisons were carried out using the CCP4 program Lsqkab (63). Subsequent multiple sequence alignments of SCO1 or COX16 with homologs were carried out using Clustal Omega and visualized using ESPript 3.x (56, 64).

The data underlying the rank plots of COX2, COX16, SCO1, SCO2, COX20 and COA6 were generated through an organelle-wide protein interaction prediction using AlphaFold Multimer v2.2 (65). We updated the genes whose UniProt IDs were missing or outdated in MitoCarta3.0 and removed unreviewed entries that could not be mapped to reviewed ones. We then removed proteins that were more than 2000 amino acids long and those that contained non-canonical amino acids. This resulted in a total of 1123 proteins whose sequences were then retrieved from UniProt. All five multimer models were run using a single random seed, and the average of all 5 ipTM scores was used as a classifier metric.

### Lipidomic analysis

Lipids from cell pellets were extracted using MTBE (Sigma Aldrich). Frozen cell pellets were first resuspended in 25 µL of PBS and 2 ul was used to normalize for protein concentration using a BCA assay (Pierce). 250 µg of protein was then resuspended in 225 µL 100% LC-grade MeOH (Fisher) containing 1 µM CoQ8 (Avanti Lipids) as an internal standard. Glass beads (100 µL; 0.5 mm; BioSpec) were added and the samples were vortexed using a Vortex Genie for 10 min (3000 rpm, 4 °C) to lyse the cells. Water was added to a total volume of 187.5 µL and then 750 µL of MTBE was added to each sample. Tubes were vortexed again for 3 min (3000 rpm, 4 °C). To separate layers, samples were centrifuged for 3 min (1000xg, 4 °C). The top, organic layer was removed into a separate microcentrifuge tube and fresh 750 µL of MTBE was added. Organic extraction was repeated, and both MTBE organic layers were pooled. Samples were lyophilized by vacuum centrifugation and resuspended in 50 µL of 20 mM ammonium acetate in 78% (v/v) MeOH, 20% IPA (Sigma Aldrich) and 2% water.

Extracted lipids were separated on an Acquity CSH C18 column (100 mm x 2.1 mm x 1.7 µm particle size; Waters) at 50 °C using the following gradient: 2% mobile phase B from 0-2 min, increased to 30% B over next 1 min, increased to 50% B over next 1 min, increased to 85% over next 14 min, increased to 99% B over next 1 min, then held at 99% B for next 7 min (400 µL/min flow rate). Column re-equilibration of 2% B for 1.75 min occurred between samples. For each analysis 1 µL/sample was injected by the autosampler. QC samples were injected at regular intervals to ensure constancy across runs. Mobile phase A consisted of 10 mM ammonium acetate (Sigma Aldrich) in 70% (v/v) ACN 30% (v/v) water with 250 µL/L acetic acid (Sigma Aldrich). Mobile phase B consisted of 10 mM ammonium acetate in 90% (v/v) IPA 10% (v/v) ACN with 250 µL/L acetic acid.

MS acquisition was performed by a Thermo Exploris 240 Orbitrap mass spectrometer. Samples were ionized by a (HESI II) source (Thermo Scientific) kept at a vaporizer temperature of 350 °C. Sheath gas was set to 50 units, auxiliary gas to 8 units, sweep gas to 1 unit. MS was operated in polarity switching mode with the spray voltage set to 3,500 V for positive mode and 2,500 V for negative mode. The inlet ion transfer tube temperature was kept at 325°C with 70% RF lens. Full MS1 scans were acquired at 22,500 resolution (at 200 m/z), max ion accumulation time of 100 ms, with a scan range of m/z 200 to 1,600. MS2 scans (Top 3) were acquired at 30,000 resolution (at 200 m/z), max ion accumulation time of 50 ms, 1.0 m/z isolation window, stepped normalized collision energy (NCE) at 20, 30, 40, and a 10.0 s dynamic exclusion. Automatic gain control (AGC) targets were set to standard mode for both MS1 and MS2 acquisitions. LC-MS files for lipidomics were processed using Compound Discoverer 3.3 (Thermo Scientific) and LipiDex 2.0. All peaks with a 1.4-23 min retention time and 100 Da to 5000 Da MS1 precursor mass were aggregated into compound groups using a 10 ppm mass tolerance and 0.4 min retention time tolerance. Peaks were excluded if peak intensity was less than 2 x 106, peak width was greater than 0.75 min, signal-to-noise ratio was less than 1.5, or intensity was < 3-fold greater than blank. MS2 spectra were searched against an in-silico generated spectral library. Spectra matches with a dot product score > 500 and reverse dot product score > 700 were retained for further analysis. Lipid MS/MS spectra that contained < 75% interference from co-eluting isobaric lipids, eluted within a 3.5 median absolute retention time deviation (M.A.D. RT) of each other and were found within at least 2 processed files were used. If individual fatty acid substituents were unresolved, then identifications were made with the sum of the fatty acid substituents. Lipid identifications were filtered with LipidDegreaser module and in-house retention time modelling. The retention time tolerance used was 0.5 min. Unreliable identifications were discarded. For BMP and PG classes, identification was performed manually in positive mode using MS2 fragments for monoacylglycerol and diacylglycerol, respectively, as has been recently reported (66). Quantification was performed only on resolved precursor peaks. Species with no MS2 fragments were identified as a separate BMP/PG class.

### Miscellaneous

Protein concentration and COX and citrate synthase activities were measured as previously described (25).

## Results

### Wild-type SCO1 and SCO2 interactomes exhibit significant overlap with respect to their composition and pathway enrichment

While SCO1 and SCO2 fulfill complementary roles in the maturation of the COX2 subassembly module and the regulation of cellular copper homeostasis (37), the suite of shared and unique protein partners that comprise their functional neighbourhoods remains largely unexplored. To address this issue, we defined the SCO1 and SCO2 interactomes using a proximity-dependent labeling strategy termed BioID, which has proven to be particularly effective for mapping protein interaction networks of various subcellular compartments including mitochondria (52, 67–69). Both C-terminally tagged SCO bait proteins were expressed at comparable levels (Figure S1A), and application of a basic filter used in a previous, large-scale study that employed SCO1 as a mitochondrial marker (54) yielded roughly 200 high-confidence prey partners per SCO bait (Supplementary Table 1). Comparative analysis showed almost complete coverage of the original dataset (54) and, critically, revealed a significant number of high-confidence prey partners specific to the SCO1 bait that were unique to our study (Figure S1B). We therefore took our high-confidence SCO1 and SCO2 prey partners and filtered them through an in-house pipeline to refine the data, and identify which interactions were enriched relative to our IMS spatial controls and occurred with proteins resident to, or with domains contained within, this mitochondrial subcompartment (Figure 1A). We then assessed the relative composition of each neighbourhood and found that SCO1 and SCO2 shared roughly 70% of potential protein partners (Figure 1B), with mitochondrial pathway (MitoPathway) enrichment analysis highlighting common enrichment for phospholipid metabolism, oxidative phosphorylation and Complex I biogenesis (Figure 1C, D). However, each SCO protein also had a significant cohort of unique neighbours (Figure 1B) and MitoPathway analysis established that SCO1 exhibited a more diverse interactome encompassing a broader range of statistically significant functional clusters when compared with SCO2 (Figure 1C, D).

**Figure 1:**
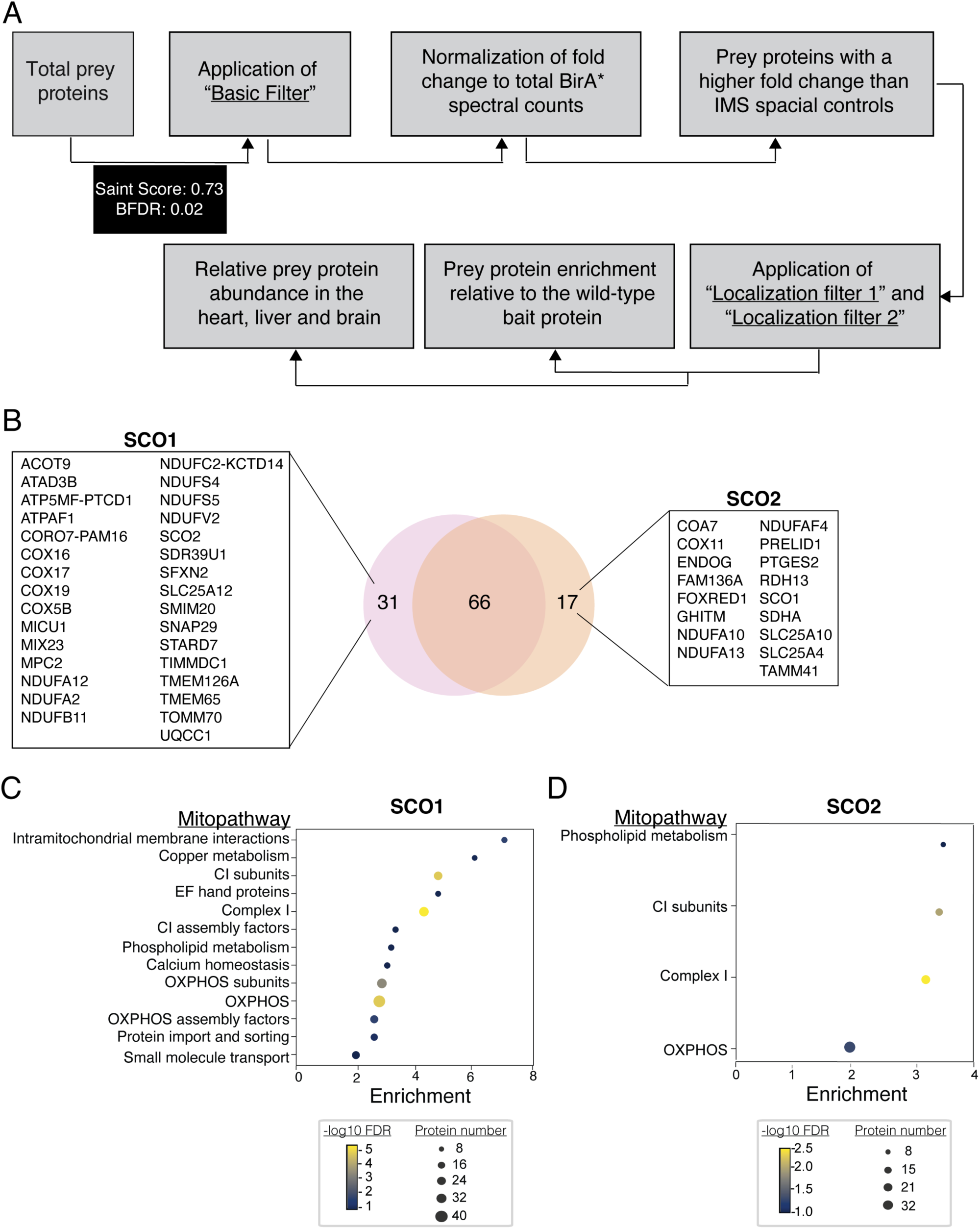
SCO1 exhibits a broader functional interactome than SCO2. *A)* Flow chart depicting the bioinformatic approach used for stepwise filtration of potential SCO interacting partners. *B)* Venn diagram comparing the wild-type SCO1 and SCO2 interactomes. *C & D)* Dot plots depicting significantly enriched MitoPathways within SCO1 and SCO2 neighbourhoods, respectively. Size of dots represents the number of identified proteins within a neighbourhood and colour represents the -log (False Discovery Rate).

### Pathogenic mutations in SCO1 and SCO2 affect their neighbourhoods both with respect to composition and enrichment for potential interacting partners

To investigate whether pathogenic *SCO* mutations affect the neighbourhoods we had mapped for the wild-type proteins, we included three SCO1 variants (SCO1 G132S, P174L and M294V) and the common SCO2 E140K variant in our proximity-ligation studies. Steady-state levels of mutant SCO baits were similar to those of their wild-type counterpart upon tetracycline induction (Figure S1A). Moreover, immunofluorescence microscopy and dot plots depicting the distribution of mitochondrial preys as a function of SAINT score indicated that all variant SCO baits were correctly localized (Figure S1C, D). We therefore applied the same data filtering pipeline (Figure 1A, Supplementary Table 1) and normalized total spectral counts for all identified prey proteins to those of the appropriate, wild-type SCO bait to quantify the impact of an allelic variant on the relative enrichment for a given protein-protein interaction (Figure 2). These analyses demonstrated that each SCO variant had a unique effect on its neighbourhood in this regard, exhibiting both increased and attenuated affinity for potential protein partners. Notably, the G132S and P174L substitutions in SCO1 and the E140K substitution in SCO2 all resulted in the detection of a small cohort of candidate interacting partners that were not identified in the wild-type background (Figure S1E).

**Figure 2:**
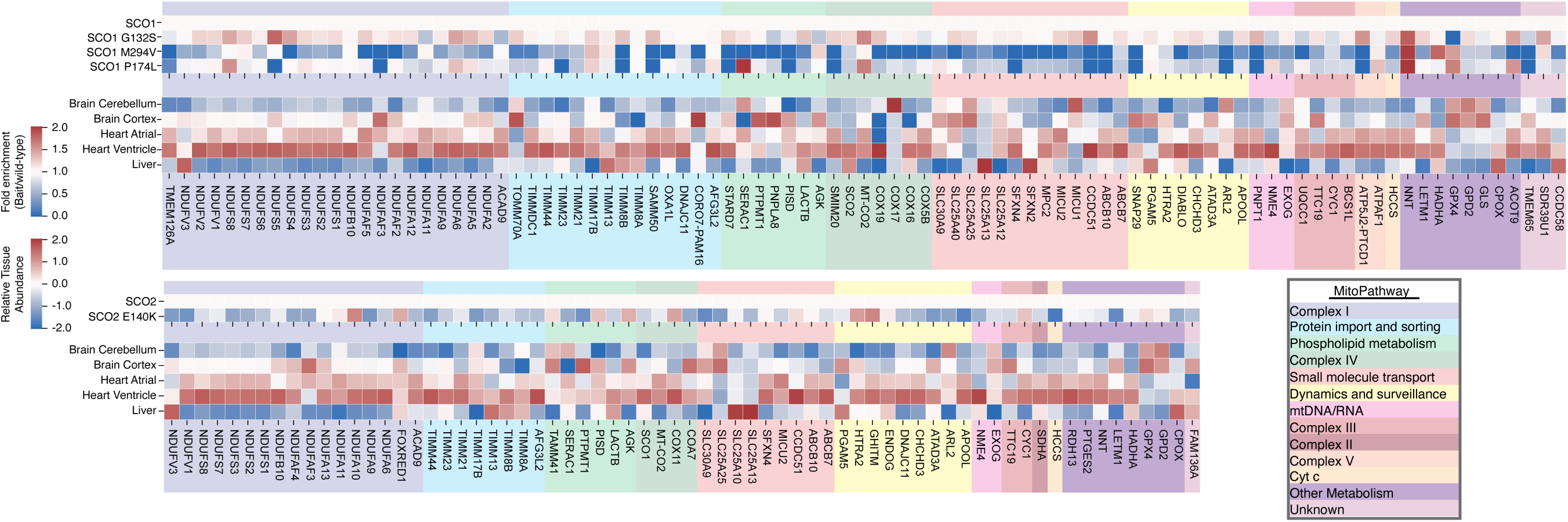
Pathogenic SCO variants exhibit an altered interactome with both decreased and increased affinity for interacting partners. Heatmaps for SCO1 and SCO2 baits (top and bottom, respectively) depicting their fold enrichment for each prey protein (upper panel) and the relative abundance of prey proteins in the heart, liver and brain (bottom panel). Prey proteins were assigned into different functional clusters and colour coded based on MitoPathway Enrichment analysis.

### The transmembrane domains of SCO1 and COX16 form hydrophobic contacts at a helix-helix interface to stabilize their interaction

Reduced affinity of a given allelic variant for an interacting partner may be more detrimental when its abundance is low, in turn offering potential insight into the tissue-specific clinical phenotypes observed across *SCO1* pedigrees and between *SCO1* and *SCO2* patients. We therefore addressed this possibility by mining the quantitative human proteome map (55) and normalizing the abundance of each prey protein in various heart, liver and brain regions to its median abundance across all measured tissues (Figure 2). Very few low abundance, candidate protein partners fit this model, which led to the identification of the SCO1 G132S-GPX4 interaction in the heart and interactions between SCO1 M294V and NME4, OXA1L and COX16 in the brain.

COX16 represents a particularly intriguing candidate given its crucial but ill-defined role in Cu_A_ site maturation (23, 70, 71), relatively low abundance in human brain (Figure 2) and reduced association with the SCO1 M294V variant (Figure 1B) linked to an isolated encephalopathy (44). Consistent with these tissue-specific expression patterns and allele-specific effects on the interaction between SCO1 and COX16, we observed that COX16 steady-state levels were relatively low in the brain of wild-type C57BL/6N mice and further reduced in a disproportionate manner upon expression of the *Sco1 M277V* allele, the murine equivalent of human SCO1 M294V (Figure 3A). Therefore, to further probe the nature of the SCO1-COX16 interaction in the context of COX assembly, we first investigated COX16 steady-state levels in wild-type (i.e. *flox*) and *Sco1* null mouse embryonic fibroblasts (MEFs). COX16 abundance was reduced roughly two-fold in the absence of SCO1, while the steady-state levels of SCO2 or COA6, two other COX assembly factors with known roles in Cu_A_ site maturation, were unaffected (Figure 3B). SCO1 and COX16 have previously been shown to physically interact (23), and *in silico* structural modeling with AlphaFold3 demonstrated that several amino acids contained within the transmembrane domain of each protein help to stabilize this interaction (Figure 3C, D). More specifically, the side chains of residues Leu98, Phe102, Leu109 and Met112 of human SCO1 and Leu25, Gly29, Phe36 and Ile39 of human COX16 form hydrophobic contacts at the a- and d-positions of a seven residue heptad repeat characteristic of a coiled-coil interface (72) that appear to be largely responsible for stabilizing this protein-protein interaction, with further stabilization and significant specificity being imparted by two salt bridges between residues Lys113 and Lys116 of SCO1 and Glu36 and Asp42 of COX16, respectively. A multiple sequence alignment of the N-terminal regions of SCO1 and SCO2 also revealed that most of these evolutionarily conserved amino acids predicted to form an interface with COX16 are unique to SCO1 when compared to SCO2 (Figure 3E). These data collectively suggest that the interaction between SCO1 and COX16 is specified and stabilized by their respective transmembrane domains.

**Figure 3:**
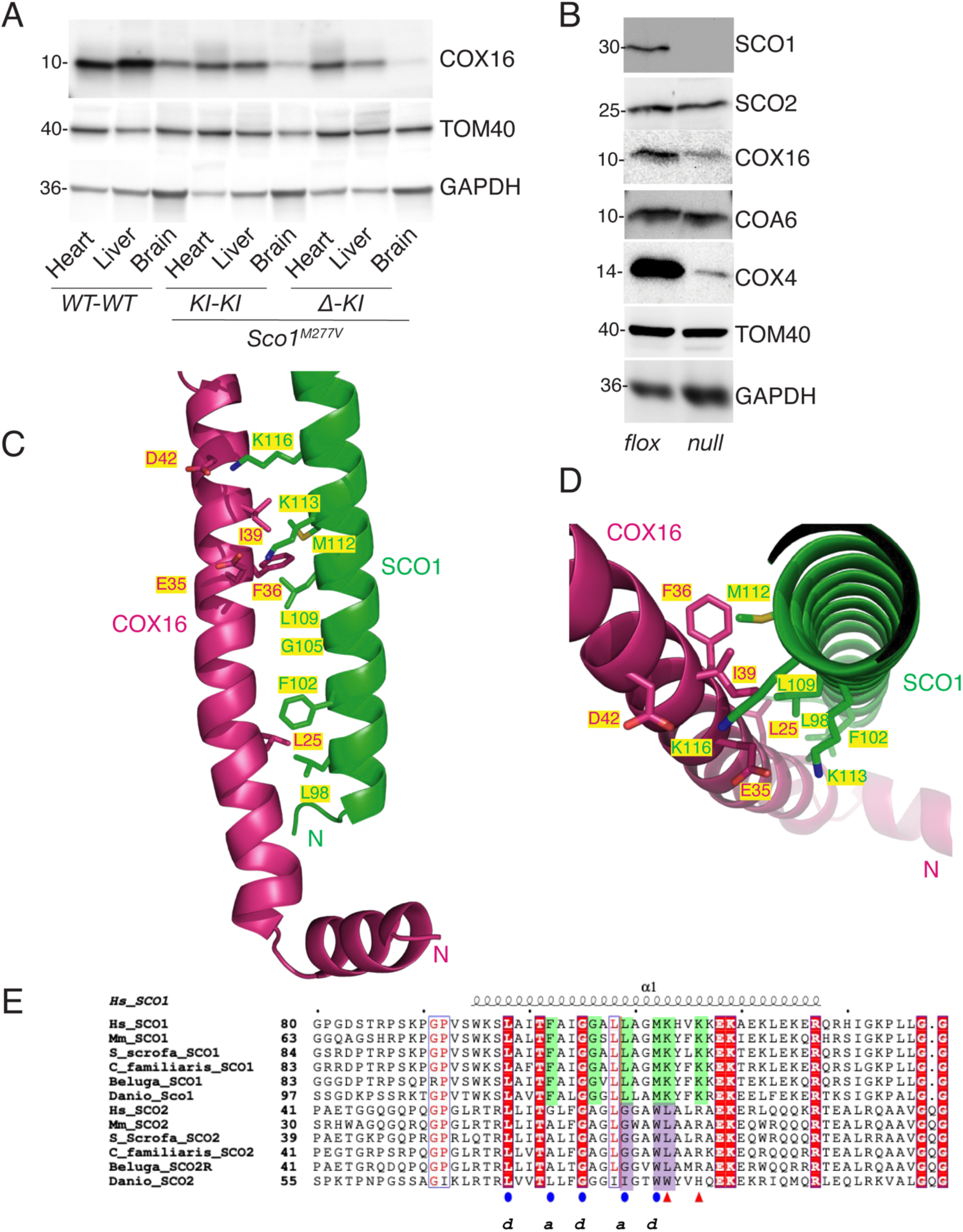
The transmembrane domains of SCO1 and COX16 stabilize their interaction. *A)* Steady-state levels of COX16 in wild-type *(WT-WT) and Sco1^M277V^ homozygous* (*KI-KI) and hemizygous (Δ-KI)* knockin mouse heart, liver and brain. *B)* SCO1, SCO2, COX16, COA6 and COX4 abundance in wild-type (*flox*) and *Sco1* knockout (*null*) MEFs. For panels *A& B)*, TOM40 and GAPDH served as mitochondrial and total loading controls, respectively. *C & D)* Ribbon representations illustrating the key residues within the N-terminal transmembrane domains of SCO1 (green) and COX16 (red) largely responsible for stabilizing the protein-protein interaction. For both proteins, amino acid side chains at the predicted interface are drawn as stick models and labeled. *E)* Multiple sequence alignment for the N-terminal regions of SCO1 and SCO2. Highly conserved residues in both SCO proteins are shaded in dark red. Residues moderately conserved between SCO1 and SCO2 are coloured in red and boxed in blue. Residues conserved only in SCO1 are shaded light green, and residues conserved only in SCO2 are shaded in lavender. Residues in SCO1 making up the proposed coiled-coil interface are marked with blue ovals, and the two lysine residues predicted to make salt bridges with COX16 residues are denoted with red triangles. Finally, residues making up the SCO1-COX16 N-helix interface are displayed and labeled in panels *C & D*.

### COX16 physically bridges SCO1 and COX2 within a ternary complex during Cu_A_ site maturation

To better understand the physical interaction between SCO1 and COX16 in the context of Cu_A_ site maturation, we used AlphaFold-Multimer to model interactions between potential partners. Pairwise predictions across the entire human mitochondrial proteome (65) using query proteins with known roles in Cu_A_ site maturation (21) suggested that COX2 may form a minimal complex with SCO1, COX16 and COX20 (Figure 4A-F). Indeed, further combinatorial analyses with AlphaFold3 using these proteins and other, candidate COX assembly factors identified a ternary complex comprised of SCO1, COX16, COX2, COX20 and COA6 (Figure 4G), which we then predicted with SCO1 and COX2 in their apo- and copper-loaded forms. SCO2 and COX17 were not included in these analyses because they require the same motif(s) as SCO1 to bind relevant partner(s) (Figures S2-S4).

**Figure 4.**
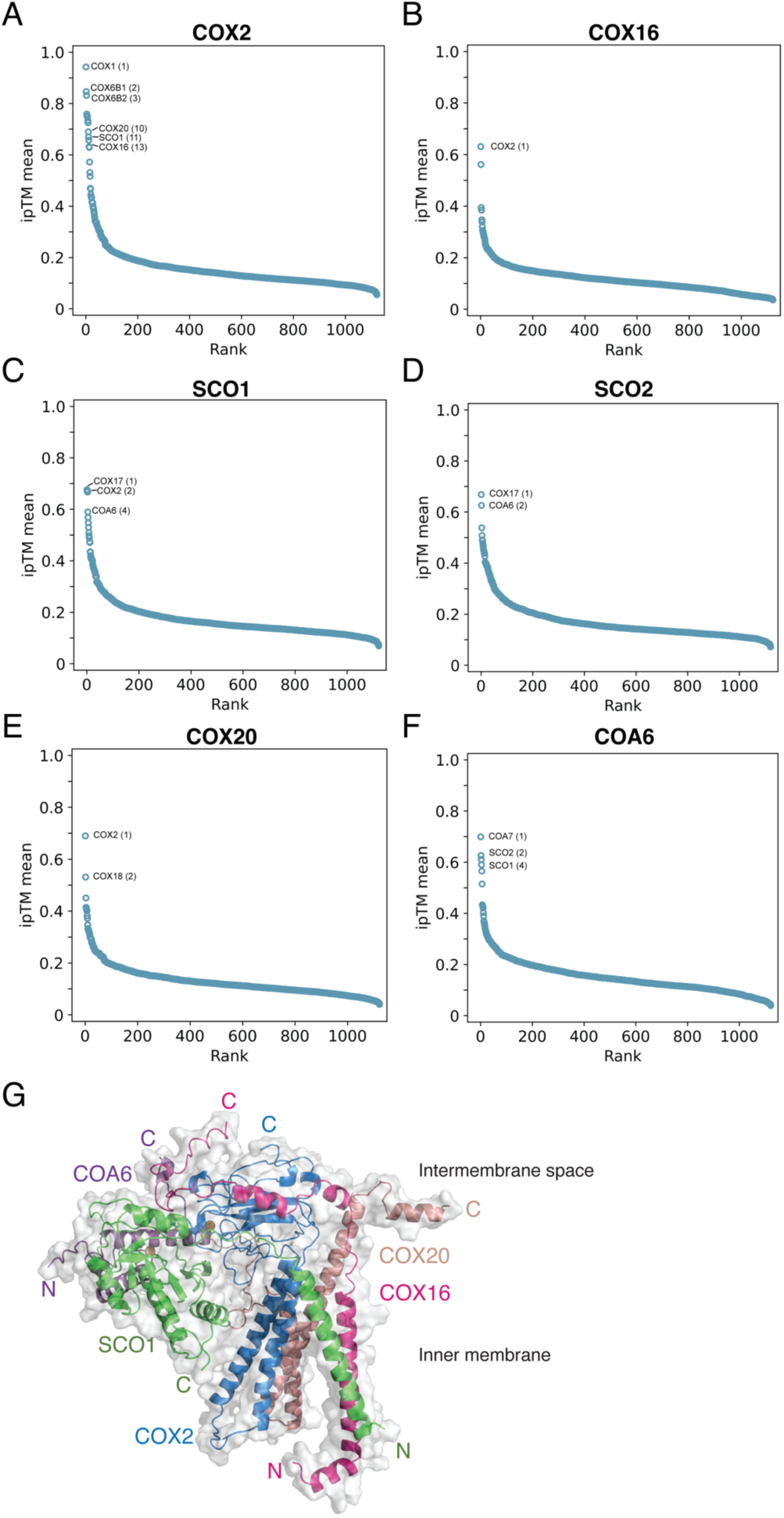
Structural modeling identifies a ternary complex comprised of SCO1, COX16, COA6, COX2 and COX20 required for Cu_A_ site maturation. *A-F)* Rank plots of the predicted interacting partners for the indicated bait proteins shown as a function of ipTM scores. *G)* AlphaFold3 model of the ternary complex responsible for copper transfer to COX2. SCO1 is coloured green, COX16 red, COA6 violet, COX2 light blue and COX20 salmon. All atomic coordinates are presented as ribbon diagrams with a semi-transparent van der Waals surface coloured in light grey. The modeled complex is viewed from within the inner membrane showing the orientations of the membrane-spanning helices. Cu^+^ ions are shown as gold spheres bound to SCO1 (one) and COX2 (two) at the known Cu^+^ binding sites for these proteins.

The modeled ternary complex indicates that SCO1 and COX16 form a highly specific heterodimer with their respective membrane spanning helices, consistent with our modeling of the two proteins in isolation (Figure 3C, D). The output Predicted Alignment Error (PAE) matrices further suggest that interactions between the membrane spanning regions of SCO1 and COX16 are most reliable, with the interactions between their water-soluble fragments becoming more reliable upon assembly of the ternary complex (Figure S5). In fact, the modeling predicts that COX16 uses its water-soluble C-terminal region to facilitate simultaneous interactions with both COX2 and SCO1 (Figure 5A, B), bridging their soluble domains and placing their respective copper-binding sites in close proximity (Figure 5C). Most soluble segment interactions between the side chains of SCO1, COX16 and COX2 involve van der Waals contacts, hydrogen bonds or salt bridges and COX16 is either in an α-helical or extended conformation relative to the known soluble domain structures of SCO1 and COX2 (Figure 5A-C). Stabilizing interactions between SCO1 and COX16 of likely importance include a salt bridge between the side chains of Asp259 of SCO1 and Arg85 of COX16, and contacts between the amide side chain of Asn80 of COX16 and the backbone amide atoms of Tyr244 of SCO1. In addition, the side chain of Trp78 of COX16 is predicted to make van der Waals contact with the polypeptide backbone atoms of residues 245 and 246 and the side chain of Arg242 of SCO1. Consistent with this prediction, the homologous Arg220 residue in yeast Sco1p (^217^KKY**<u>R</u>**VYF^223^) has previously been shown to be critical for its interaction with Cox2p (73). This conserved motif spans the C-terminus of the predicted helix α4 and the subsequent strand β6 in human SCO1 (^239^RAY**<u>R</u>**VYY^245^) and resides at the SCO1:COX16 soluble domain interface (Figure 5A). Finally, a predicted salt bridge between Asp76 of COX16 and Arg239 of SCO1 along with likely contacts between Phe75 of COX16 and Leu135 of SCO1 are predicted to assist in the stabilization of this protein-protein interaction (Figure 5A). Predicted interactions between the soluble segments of COX16 and COX2 are more extensive and likely include a salt bridge between Glu66 of COX16 and Lys98 of COX2 along with several potential van der Waals contacts including Leu52 and Leu56 of COX16 and Leu216 of COX2, Ile61 of COX16 and Met153 of COX2, and Ile70 of COX16 and Leu179 of COX2 (Figure 5B).

**Figure 5.**
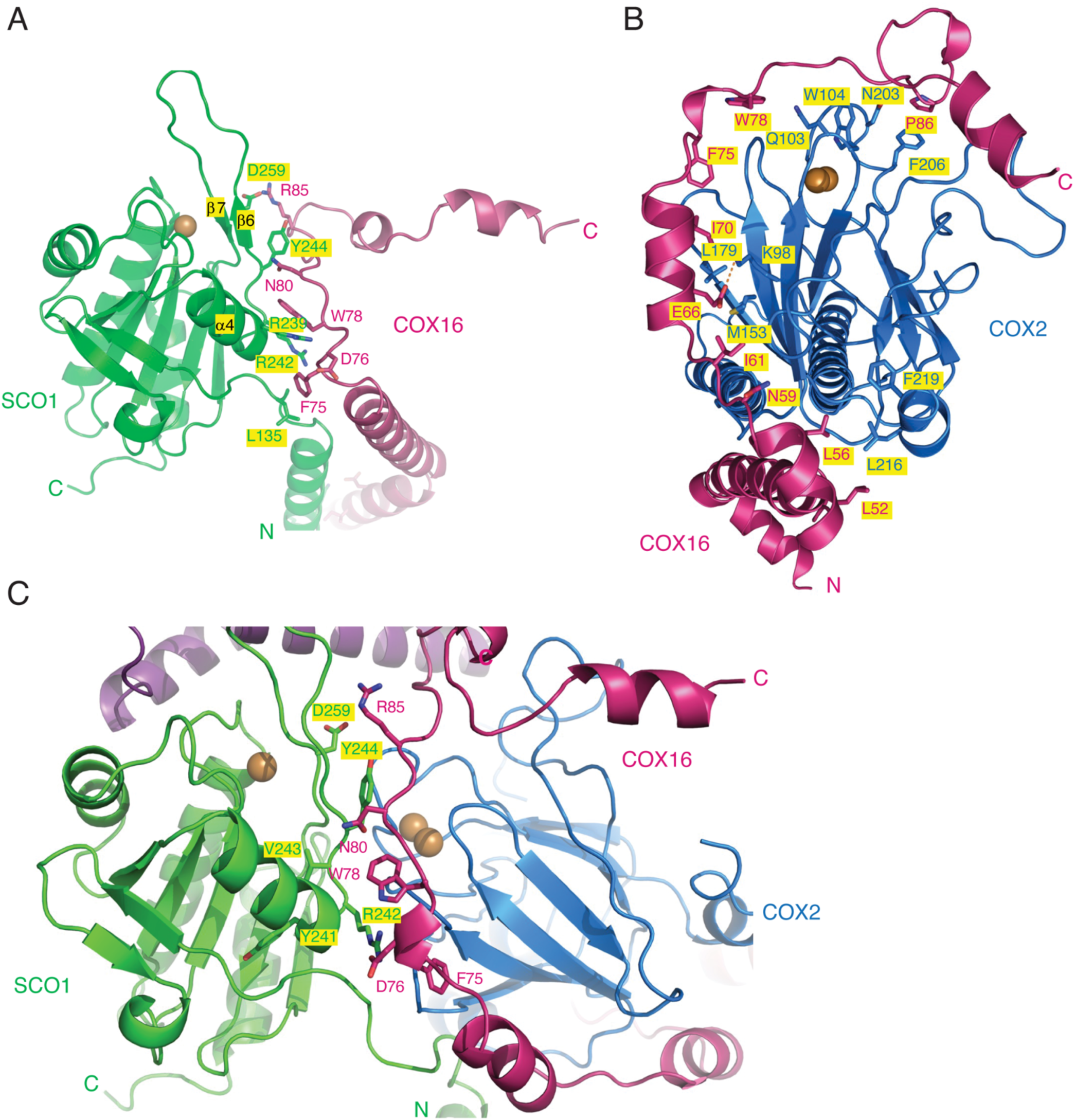
Predicted SCO1:COX16 and SCO1:COX2 interactions within the ternary complex with a closeup view of the interface. *A)* Here only SCO1 and COX16 are shown for the modeled ternary complex (see Figure 3G), with their N-terminal membrane-spanning helices on the lower right and the Cu^+^ binding site of SCO1 at the upper left. The carboxylate side chain of SCO1 residue Asp259, adjacent to the Cu^+^ ligand His260, makes a salt bridge with the side chain of Arg85 of COX16. Again, Cu^+^ is depicted as a gold sphere. Side chains on COX16 and SCO1 that are predicted to be in van der Waals contact or make hydrogen bonds at the predicted interface are shown as stick models and are coloured and labeled appropriately. *B)* A similar view of the COX16:COX2 subcomplex with COX16 residues that are predicted to interact with residues on COX2 shown as stick models and coloured red and blue, respectively, and labeled accordingly. *C)* A combined view including regions of SCO1, COX16 and COX2 at the soluble domain interface, drawn as in panels *A)* and *B)*. The orientation of the molecular subcomplex is similar to panel *A)*, and side chains likely interacting with neighbouring molecules are coloured appropriately and labeled as in panels *A & B)*. For panels *A-C)*, models are viewed approximately from the intermembrane space above the presumed plane of the mitochondrial inner membrane.

Gln103, Trp104 and Asn203 of COX2 are also predicted to interact with side chains of amino acids from COX16 and SCO1 (Figure 5B, C), and the relative positioning of their side chains, especially residues involved in salt bridges, appears to depend on the metalation state of SCO1. Prior to copper incorporation into the Cu_A_ site, its apo-state alters the interface between COX2, COX16 and SCO1, especially for Gln103 and Trp104 of COX2 and nearby residues 80-84 of COX16 (Figure S6). For SCO1, the absence of bound copper very likely impacts the local conformation of residues Asp259 and Asp171 which normally would form salt bridges with the side chains of Arg85 on Cox16 and Arg25 and Arg64 on COA6, respectively. Taken together, our modeling of the ternary complex critical to Cu_A_ site maturation suggests that these non-covalent intermolecular interactions facilitate the precise association of SCO1 within the ternary complex and communicate its copper bound state to COA6 and COX16 and, by extension, COX2.

### COX16 abundance is reduced upon impairment of COX2 subassembly module maturation

Our data strongly suggest that SCO1 is required for the stability of COX16 which in turn serves as a protein bridge within a ternary complex to spatially align the copper-binding sites of SCO1 and COX2 to facilitate Cu_A_ site maturation. To further evaluate the importance of the interaction between COX16 and SCO1, we assessed COX16 steady-state levels in wild-type or *Sco1* null MEFs overexpressing a cDNA encoding the murine equivalent of each pathogenic SCO1 variant or the wild-type protein. Overexpression of wild-type SCO1 or SCO1 M277V normalized the abundance of COX16 and COX4 (Figure 6A), an established marker of holoenzyme levels in a murine background (49, 74, 75). In contrast, the steady-state levels of COX16 were unaffected by overexpressing the SCO1 G115S or P157L variants (Figure 6A), which have previously been shown to have very little residual function as COX assembly factors (50). To determine if our MEF findings were recapitulated in human cells, we next detected COX16 abundance in *SCO1* patient fibroblasts harboring the *P174L* or *M294V* alleles. Endogenous expression of either allelic variant resulted in a roughly two-fold reduction in COX16 abundance when compared to control fibroblasts (Figure 6B).

**Figure 6.**
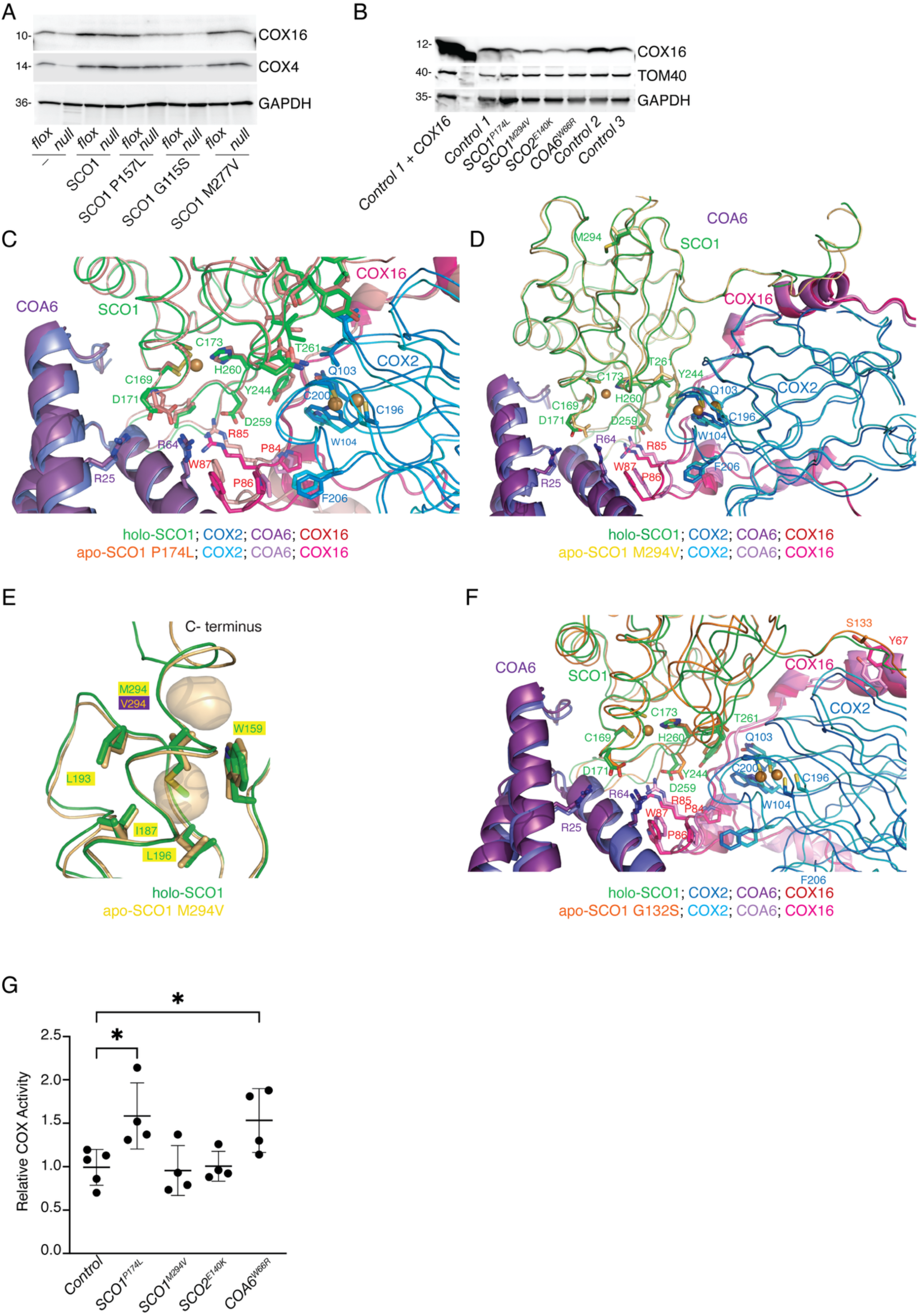
Impact of allelic variants on the ternary complex and COX16 stability in the context of Cu_A_ site maturation. *A)* Steady-state levels of COX16 and COX4 in wild-type (*flox*) and *Sco1* knockout (*null*) MEFs alone (-) or overexpressing a given SCO1 allelic variant or the wild-type protein. GAPDH served as a loading control. *B)* COX16 abundance in *Control* fibroblasts and fibroblasts isolated from patients with mutations in *SCO1*, *SCO2* or *COA6*. A control lysate overexpressing COX16 was included as a positive control. TOM40 and GAPDH served as mitochondrial and total loading controls, respectively. *C)* View of wild-type Cu^+^-loaded human SCO1 (holo-SCO1; green) in complex with COX16 (dark red), Cu^+^-loaded COX2 (dark blue), and COA6 (dark purple) overlaid on the complex of apo-SCO1 P174L (tawny), COX16 (pink-red), apo-COX2 (light blue) and COA6 (lavender). *D)* as in panel *C)* but with apo-SCO1 M294V. *E)* Close up view of *D)* illustrating the internal voids created by the M294V substitution shown as pale gold transparent surfaces. *F)* as in panel *C)* but with apo-SCO1 G132S. For panels *C)*, *D)* and *F)* **t**he Cu^+^ ions are shown as bronze spheres. Side chains involved in interactions between the various subunits are shown as stick models with carbons coloured appropriately. View is approximately from the presumed inner membrane towards the intermembrane space with SCO1 in the foreground and COX16 in the background. Rotated approximately 90 degrees counterclockwise with respect to Figure S5. *G)* COX activity in *Control*, *SCO1*, *SCO2* and *COA6* patient fibroblasts overexpressing COX16 normalized to that of the untransduced, parental line (*Controls*, n=5; patient backgrounds, n=4). *p<0.05 detected by a one-way ANOVA paired with Dunnett’s post hoc test.

To further probe the structural basis of our molecular observations, we modeled the potential impact of all three SCO1 variants on the ternary complex and found that each mutant likely presents a unique deficiency with respect to maturation of the COX2 subassembly module. In agreement with previous NMR and molecular studies of soluble SCO1 truncates (76, 77), the Pro174 to Leu substitution induces small, localized changes to the side chain positions of the copper ligand residues Cys169 and Cys173, leading to a closer sulphur-sulphur distance and subsequent disulphide bond formation (Figure 6C). These changes also impact the positions of the carboxylate groups of Asp171 and Asp259 which then likely relay the redox state of the CxxxC motif of SCO1 and the lack of copper to COA6 and COX16, respectively. Moreover, COX2 likely senses the CxxxC redox state of SCO1 P174L via altered interactions with COX16 in the vicinity of Arg85 and other nearby residues (Figure 6C). In contrast, the Met294 to Val substitution modestly alters the position of the α-helix adjacent to the copper-binding helix (Figure 6D); however, more importantly, it introduces internal cavities in the SCO1 structure near the C-terminus (Figure 6E) which reduce the number of side chain-side chain van der Waals contacts, increase overall mobility and in turn reduce the thermal stability of the protein. Consistent with this idea, endogenous SCO1 M294V is virtually undetectable in patient fibroblasts (44). The Met294 to Val mutation is also likely to directly alter the position of the C-terminus of the Cu-binding helix of SCO1 as the Val side chain will introduce a van der Waals repulsion with the side chains of Leu193 and Ile187, inducing small tertiary changes that again result in modest structural perturbations in the positions of copper ligands Cys169 and Cys173 (Figure 6D). Finally, for the SCO1 G132S mutant, the Gly to Ser substitution changes the orientation of the SCO1 N-terminal transmembrane helix (Figure 6F), perturbing its interactions with COX16 and in turn destabilizing the SCO1-COX2 interaction.

Consistent with the idea that the stability of the ternary complex is impacted by more than the SCO1-COX16 interaction, we found that pathogenic mutations in other gene products known to play a role in Cu_A_ site maturation also led to a reduction in COX16 abundance (Figure 6B). To further investigate the functional relevance of COX16 in this context, we overexpressed a *COX16* cDNA in control, *SCO1*, *SCO2* and *COA6* patient fibroblasts and measured COX activity in each background. COX16 overexpression was only able to partially complement the isolated COX deficiency in *SCO1* patient fibroblasts harboring the *P174L* allele and in *COA6* patient cells (Figure 6G). This finding agrees with our molecular and *in silico* analyses which collectively indicate that COX16 does not interact with SCO2 and that its association with SCO1 is severely attenuated by the Met294 to Val substitution.

### SCO1 and SCO2 alter phospholipid metabolism in an allele-specific manner

While the bulk of mitochondrial phospholipids are synthesized within the endoplasmic reticulum and imported into mitochondria, the organelle contains dedicated machinery to synthesize multiple classes of membrane phospholipids (78). BioID for both SCO1 and SCO2 identified a diverse subset of well-defined mitochondrial phospholipid biosynthetic enzymes, including those involved in the biosynthesis of phosphatidylethanolamine (PE; PISD), phosphatidic acid (PA; AGK), and phosphatidylglycerol (PG) and cardiolipin (CL; TAMM41, PTPMT1). Additionally, we identified potential interactions with the poorly characterized lipases LACTB, SERAC1 and PNPLA8 which have been linked to mitochondrial phospholipid metabolism but have ill-defined biological functions (79–81) (Figure 2).

The allele-specific enrichment of interactions with phospholipid-related enzymes led us to explore if loss of SCO1 or SCO2 function leads to unique changes to the cellular lipidome. To test this idea, we performed untargeted lipidomics using fibroblasts from two control, two *SCO1* patient (*SCO1^P174L^, SCO1^M294V^*) and two *SCO2* (*SCO2^E140K/Δ^, SCO2^E140K/R171W^*) patient backgrounds. Principal component analysis from the lipidomic measurements illustrated that replicates from *SCO1^P174L^* patient fibroblasts clustered separately from controls and *SCO1^M294V^* fibroblasts in the first and second principal components, confirming global cellular lipid alterations unique to that background. In contrast, the *SCO2* patient backgrounds were clustered together more tightly but remained separate from the control fibroblast lines (Figure 7A). To further probe these observations, we examined each patient lipidome in detail and found significant increases in Bis(monoacylglycerol)phosphate (BMP) or BMP/PG lipid species that were unique to the *SCO1^P174L^* patient background and one of the two *SCO2* patient backgrounds (*SCO2^E140K/Δ^*, Figure 7B-D). Overexpression of the appropriate wild-type *SCO* cDNA partially rescued the high abundance of these lipid classes in both patient backgrounds, although not all reached statistical significance (Figure 7E, F). Critically, we did not observe drastic changes in the overall abundance of other mitochondrial phospholipids such as CL (Figure S7A) and neither BMP nor PG abundance was increased in separate experiments examining other COX-deficient patient fibroblast lines with mutations in *COX10* or *SURF1* (Figure S7B, C). Taken together, these results corroborate the BioID data and further suggest that SCO1 and SCO2 impinge upon phospholipid metabolism.

**Figure 7.**
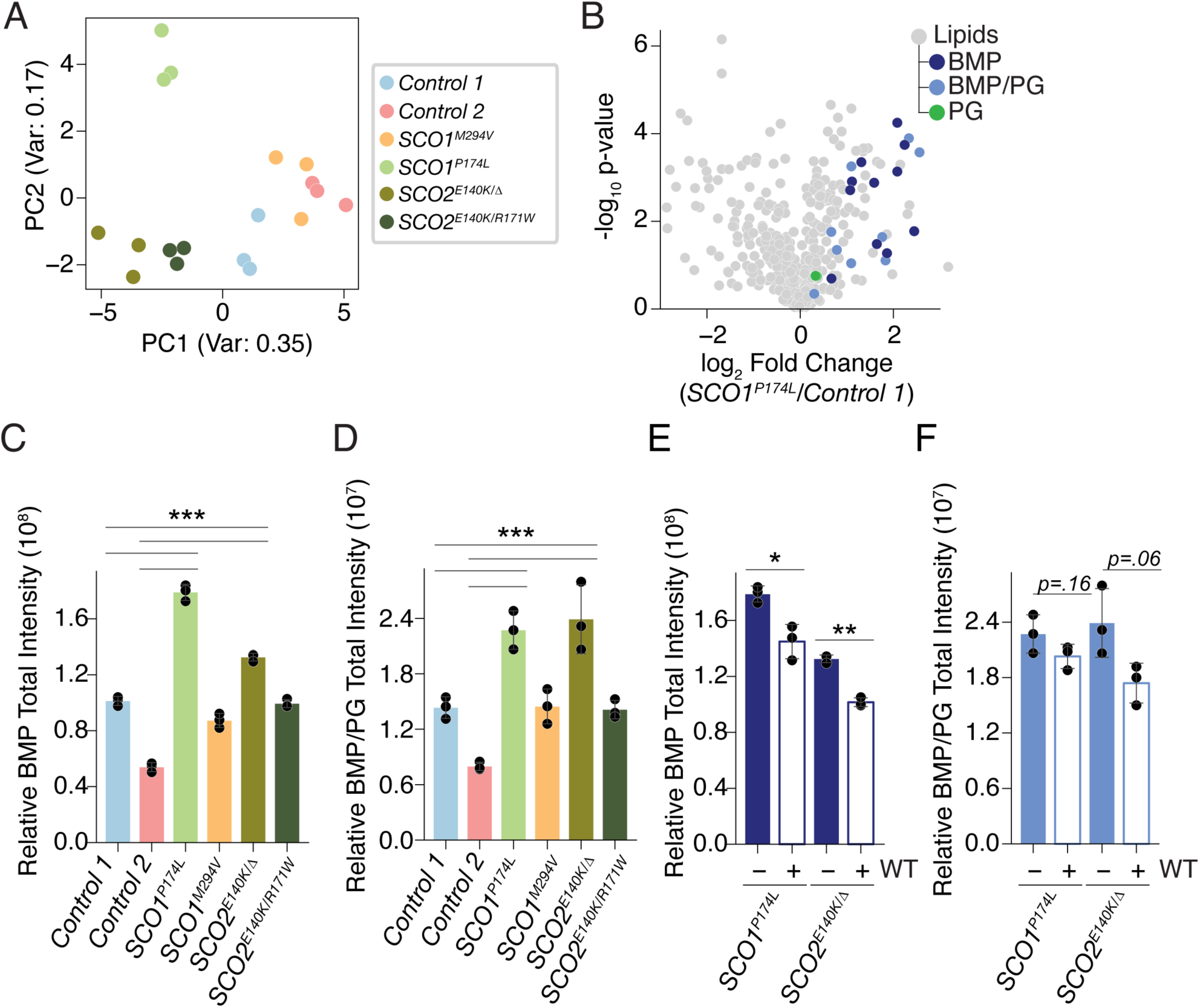
*SCO1* and *SCO2* mutations disrupt cellular phospholipid metabolism in an allele-specific manner. *A)* Principal component analysis of lipidomics data collected from *SCO1* patient (P174L and M294V), *SCO2* patient (E140K/Δ, E140K/R171W) and *Control* (MCH46 [*Control 1*] and MCH58 [*Control 2*]) fibroblast lines. *B)* Relative lipid abundance in *SCO1 ^P174L^*patient fibroblasts compared to *Control 1* fibroblasts, versus statistical significance. Identified Bis(monoacylglycero)phosphate (BMP) and Phospatidylglycerol (PG) species are coloured in dark blue and green, respectively. Lipids that could only be identified as being either BMP or PG (BMP/PG) are coloured in light blue. *C)* Combined relative intensities of all detected BMP species in *SCO1* patient (P174L and M294V), *SCO2* patient (E140K/Δ, E140K/R171W), and *Control* fibroblast lines. *E)* Combined relative intensities of all detected BMP/PG species in *SCO1* patient (P174L and M294V), *SCO2* patient ((E140K/Δ, E140K/R171W), and *Control* fibroblast lines. *E)* Combined relative intensities of all detected BMP species in *SCO1^P174L^*and *SCO2^E140K/Δ^* patient fibroblasts alone (-) or overexpressing wild-type SCO1 or SCO2 (WT), respectively. *F)* Combined relative intensities of all detected BMP/PG species in *SCO1^P174L^* and *SCO2^E140K/Δ^* patient fibroblasts alone (-) or overexpressing wild-type SCO1 or SCO2 (WT), respectively. For all panels, n=3 for each cell line or condition.

## Discussion

Identifying mechanisms that underlie the tremendous clinical heterogeneity tied to mitochondrial disorders remains a challenge with respect to their diagnosis and the rational design of therapeutics.

Here, we used BioID to characterize the SCO1 and SCO2 neighbourhoods and explore how their respective interactomes were perturbed by known pathogenic point mutants. We observed allele-specific effects on the affinity of each SCO protein for known or potential partners that disrupt interactions with proteins in both canonical and non-canonical pathways, and follow-up studies shed potential mechanistic insight into how this might contribute to the distinct clinical presentations observed across *SCO1* pedigrees and between *SCO1* and *SCO2* patients.

SCO1 and SCO2 fulfill independent, cooperative roles in COX2 maturation during holoenzyme assembly that depend on their ability to bind copper (32), and our BioID data are consistent with a two-step process in which SCO2 acts upstream of SCO1 to stabilize the nascent COX2 polypeptide chain (25, 33). A previous study argued that SCO2 is a substrate of COA7 (24), and we observed that their association is significantly impaired by the SCO2 E140K substitution. Given that the SCO2 E140K fusion protein is also less able to biotinylate COX2, it is tempting to speculate that the thiol reductase activity of COA7 is required for SCO2 to interact with the nascent polypeptide chain so that it can contribute to Cu_A_ site reduction to accommodate copper loading by SCO1. While SCO1 and SCO2 are clearly in close physical proximity based on their ability to reciprocally label one another, our modeling data emphasize that they do not form a complex and that these closely related paralogs utilize the same, shared secondary structural elements and molecular surfaces to interact with COX17 and COX2 (73). In fact, although other interpretations of our data are possible, we favour the idea that COA6 and SCO2 act as thiol reductases within the same complex to prime the Cu_A_ site for metalation and that SCO2 is subsequently displaced to allow for the assembly of a ternary complex comprised of SCO1, COX16, COX20, COA6 and COX2. The presence of COA6 in both complexes may reflect the fact that it is required to stabilize the redox state of the cysteines within the primed apo-Cu_A_ site prior to copper transfer from SCO1. Consistent with this overarching idea, COA6 binds both SCO proteins but has been shown to preferentially interact with SCO1 (28). We suspect that the inability to detect significant COA6 biotinylation upon expression of either SCO bait protein likely reflects the relative positioning of the biotin ligase within each ternary complex. Why expression of the SCO2 bait did not yield COX17 biotinylation above background levels is unclear; however, it may suggest that SCO2 binds copper constitutively to fulfill its redox function (34) and thus, unlike SCO1, does not require repeated rounds of interaction with COX17.

While it is clear SCO1 acts downstream of SCO2 as a copper insertase (34), the protein constituents of the associated ternary complex, their respective roles during copper transfer to COX2 and the impact of pathogenic mutations on this biochemical process and tissue-specific disease etiology have not been fully explored. Our study sheds some light on this multi-layered issue. Although both SCO proteins can be modeled to form a ternary complex with COA6 and COX17, we were unable to incorporate either SCO2 or COX17 within the ternary complex containing SCO1, COX16, COX2, COA6 and COX20. This incompatibility is likely explained by the fact that both human SCO1 (^239^RA<u>YRVY</u>Y^245^) and SCO2 (^203^HS<u>YRVY</u>Y^209^) use the same, evolutionarily conserved YRVY motif to interact with both COX17 and COX2 (73). Thus, SCO1 must be copper-loaded by COX17 either prior to or upon assimilating into the ternary complex that facilitates Cu_A_ site maturation. Critically, Arg242 and Tyr244 of SCO1 both reside at the interface with COX17 and COX16, which serves as an essential bridge between SCO1 and COX2. A role for COX16 in this context has previously been proposed based on the findings of physical pulldown studies in wild-type and *COX16* knockout cells (23), and our structural modeling highlights that the necessary residues for this bridging function are lacking in the pathogenic COX16 R82* truncation mutant (71), which serves to impair the positioning of the copper-binding sites of SCO1 and COX2 and hamper efficient ligand exchange. Strikingly, all three pathogenic SCO1 variants label COX2 to a greater extent than the wild-type protein, indicating that the residency time is increased owing to the inability of each mutant to efficiently form a productive ternary complex and catalyze copper transfer to the nascent COX2 polypeptide chain. This observation implies that timely dissociation of the SCO1:COX16:COX2 complex requires a conformational change tied to efficient copper transfer from SCO1 to COX2. Consistent with this idea, our BioID data demonstrate a dampened ability of all allelic SCO1 variants to interact with COX17, and reduced association of SCO1 P174L and SCO1 M294V for COX16. COX16 and SCO1 abundance is low in the brain relative to the heart and liver, and the M294V substitution destabilizes the tertiary fold of SCO1. Our findings therefore lead us to propose that the reduced COX16 expression we and others (70) have observed in the brain exacerbates the loss of SCO1 function attributable to the M294V substitution analogous to a synthetic sick interaction observed in classical genetics studies, thereby explaining the unique tissue-specific clinical course of disease in this *SCO1* pedigree.

Unexpectedly, our BioID data identified several interactions between SCO1 and SCO2 and proteins involved in cellular phospholipid metabolism that were uniquely perturbed in an allele-specific manner and may therefore contribute to the clinical heterogeneity observed between *SCO* pedigrees. These observations align with complexome profiling results in yeast, which identified large molecular weight complexes containing mitochondrial phospholipid biosynthetic enzymes and ancillary factors required for the assembly of OXPHOS complexes, including Sco1p and Sco2p (82). While the exact biological function of these complexes is unclear, their existence suggests there is an underappreciated link between COX assembly factors, holoenzyme biogenesis and mitochondrial phospholipid biosynthesis that is worthy of further exploration. Importantly, our untargeted lipidomics experiments found that phospholipid metabolism in *SCO1* and *SCO2* patient fibroblasts was uniquely altered when compared to *COX10* or *SURF1* patient backgrounds, emphasizing that our findings were not attributable to the isolated COX deficiency *per se* and further suggesting that SCO1 and SCO2 may have undiscovered roles in these metabolic pathways. Perhaps surprisingly, we did not observe decreases in the abundance or changes to the composition of mature cardiolipin in the *SCO1* and *SCO2* patient backgrounds, a phenomenon that has previously been described in cell culture models with dysfunctional mitochondria (83, 84). We instead observed increases in the abundance of lipids identified as either BMP or PG. Many of the mechanisms of BMP biosynthesis have only recently begun to be elucidated (85, 86), and how PG is exported from mitochondria and potentially remodelled for BMP biosynthesis is not well understood. Interestingly, the interaction between SCO1 and SERAC1 was uniquely increased by the P174L substitution, and mutations in *SERAC1* have already been linked to altered PG remodelling (81) with *SERAC1* patients presenting with MEGD(H)EL syndrome with varying hepatic involvement (87, 88) while the *SCO1 P174L* patient succumbed from a neonatal hepatopathy (43). However, our results suggest that SCO1 and SCO2 may both play a role in PG and BMP metabolism and investigating these mechanisms is an area of future investigation.

In summary, we used BioID to characterize the neighbourhoods of wild-type and pathogenic variants of SCO1 and SCO2 to garner insight into the clinical heterogeneity of the associated diseases and mitochondrial disorders in general. We found that the wild-type interactomes were enriched for fairly diverse functional clusters and captured potential partners with roles in both canonical and non-canonical biochemical pathways, which was consistent with a recent study that systematically mapped the human mitochondrial complexome in HEK293T cells (bioRxiv 2026.05.29.728872). We then used a number of follow-up approaches to identify a potential mechanism underlying tissue-specific disease tied to COX assembly, a canonical pathway, and cellular phospholipid metabolism, a non-canonical pathway. Collectively, our data emphasize the potential of proximity ligation to further define the molecular roles of disease-causing variants that perturb mitochondrial function and suggest that SCO proteins impinge upon cellular phospholipid metabolism.

## Acknowledgements

The authors are indebted to Dr. George Katselis (University of Saskatchewan) for preliminary mass spectrometry analyses and Ms. Nikki Forsberg (University of Saskatchewan) for identification of the linker sequence used between the bait and biotin ligase. We would also like to acknowledge Drs. Maina Kava (Perth Children’s Hospital, Australia) and Lawrence Greed (PathWest Laboratory Medicine, Australia) along with the affected family for consenting to send us *COX16* patient fibroblasts. The research reported in this publication was supported by grants-in-aid of research from the Canadian Institutes of Health Research (CR3-184665, S.C.L.), the National Institute of General Medical Sciences of the National Institutes of Health (R35GM152102, V.M.G.; R35GM131795, D.J.P.), the Robert A. Welch Foundation (A-2280-20260402, V.M.G) and the BJC Investigator program (D.J.P.). D.J.P. is an investigator of the Howard Hughes Medical Institute. The content is solely the responsibility of the authors and does not necessarily represent the official views of the NIH.

## Author Contributions

S.C.L. and S.G. conceived the project. S.G., Z.N.B., R.G., A.B.S. and S.M. collected and analyzed the data. H.A. helped with .mzxml file conversion and subsequent processing and analysis of BioID data. S.C.L., S.G., S.M., Z.N.B., D.P. and V.M.G. drafted and edited the manuscript.

