## Supplementary tables and figures for "Pathogenic variants of the mitochondrial copper chaperones SCO1 and SCO2 reshape their respective interactomes and disrupt cellular phospholipid metabolism"

**2 Supplementary Tables, 5 Supplementary Figures**

**Supplementary Table 1.** Total number of preys per bait protein upon sequential data filtration.

| Baits / Filters | SCO1 WT-BirA* | SCO1 PL-BirA* | SCO1 GS-BirA* | SCO1 MV-BirA* | SCO2 WT-BirA* | SCO2 EK-BirA* |
|---|---|---|---|---|---|---|
| Total Prey Proteins | 1109 | 1312 | 1074 | 1213 | 1037 | 930 |
| Basic Filter | 266 | 204 | 266 | 171 | 214 | 197 |
| Localization filter 1 | 194 | 149 | 194 | 128 | 155 | 149 |
| Localization filter 2 | 97 | 83 | 102 | 61 | 83 | 80 |

**Supplementary Table 2.** List of antibodies

| Antibodies | Source | Identifier | Dilution | Type |
|---|---|---|---|---|
| SCO1 | In house | (1) | 1:1000 | Rabbit, polyclonal |
| SCO2 | In house | This study[1] | 1:1000 | Rabbit, polyclonal |
| COX4 | Abcam | ab110272 | 1:1000 | Mouse, monoclonal |
| FLAG | Sigma | F1804 | 1:1000 | Mouse, monoclonal (western blot) |
| FLAG | Sigma | F7425 | 1:300 | Rabbit, polyclonal (immunofluorescence) |
| HSPA9 | Invitrogen | MA3-028 | 1:300 | Mouse, monoclonal |
| TOM40 | Proteintech | 18409-1-AP | 1:5000 | Rabbit, Polyclonal |
| COX16 | Proteintech | 19425-1-AP | 1:1000 | Rabbit, Polyclonal |
| COA6 | Proteintech | 24209-1-AP | 1:1000 | Rabbit Polyclonal |
| GAPDH | BioRad | 12004168 | 1:5000 | hFAB, Rhodamine conjugated monoclonal |
| anti-Mouse IgG (H+L) secondary | BioRad | 170-6516 | 1:5000 | Goat, HRP conjugated polyclonal |
| anti-Rabbit IgG (H+L) secondary | BioRad | 170-6515 | 1:5000 | Goat, HRP conjugated polyclonal |
| anti-Mouse IgG (H+L) secondary, cross-adsorbed | Invitrogen | A-11005 | 1:200 | Goat, Alexa Fluor 594 conjugated polyclonal |
| anti-Rabbit IgG (H+L) secondary, highly cross-adsorbed | Invitrogen | A-21206 | 1:200 | Donkey, Alexa Fluor 488 conjugated polyclonal |

[1]A human SCO2 antibody exhibiting cross-reactivity with the murine ortholog was generated by injecting rabbits with a synthetic peptide (CRTEALRQAAVGQGDFHLLDH). Peptide conjugation and emulsification along with purification of the crude polyclonal antiserum was done as previously described in (2).

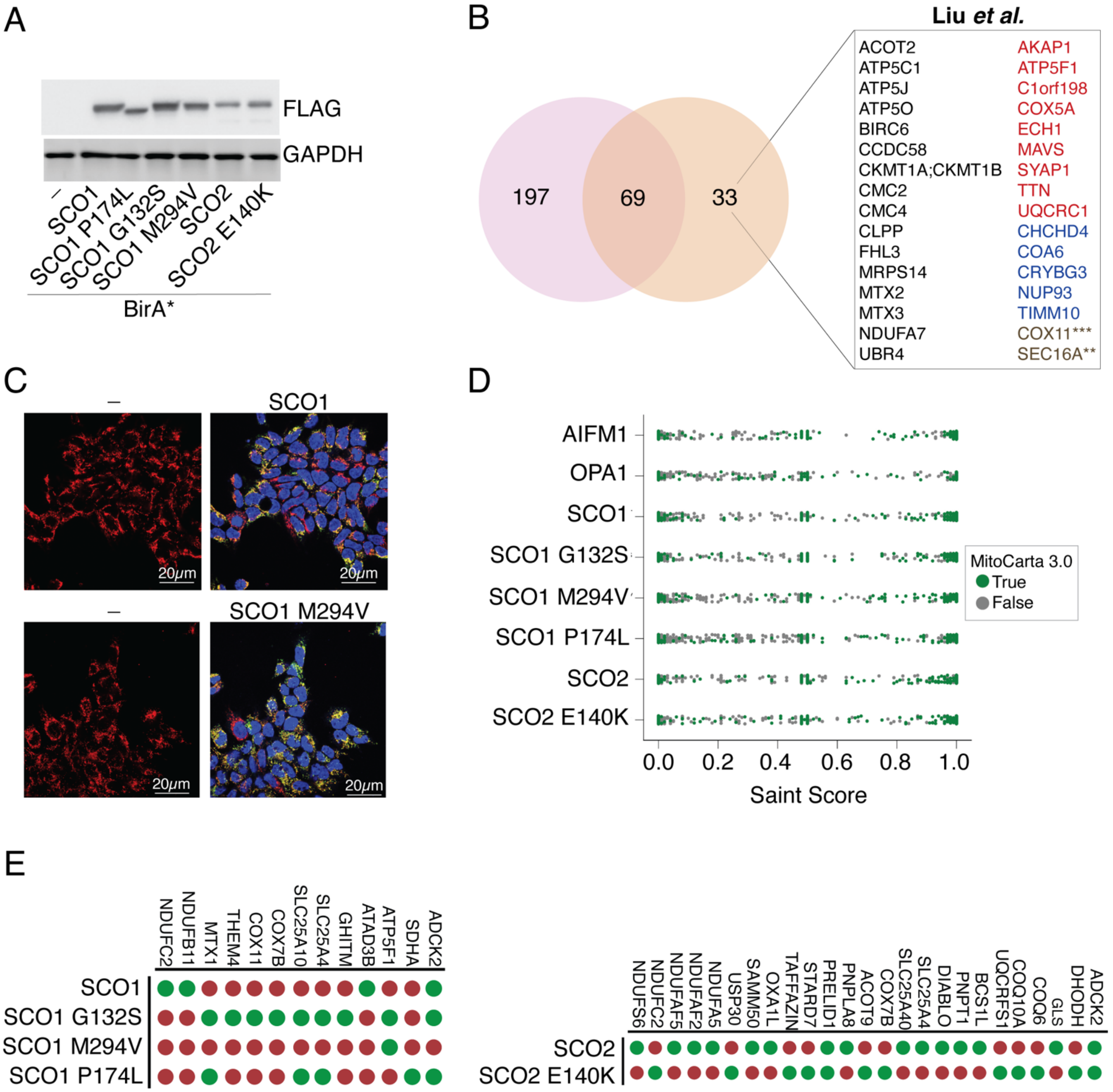


Figure S1

**Figure S1. SCO bait proteins are expressed at comparable levels and localize to mitochondria.** *A)* Steady-state levels of C-terminally FLAG tagged SCO1 and SCO2 bait proteins detected with a FLAG antibody. GAPDH served as a loading control. *B)* Venn diagram illustrating the overlap between the wild-type SCO1 interactomes in our study and that of Liu *et al.* (3). Black font indicates unique prey proteins, red font signifies prey proteins eliminated in our study by the basic filter, blue font denotes prey proteins with higher spectral counts in the spatial controls and brown font highlights prey proteins only detected by allelic SCO1 variants (asterisks indicate how many SCO1 variants interacted with a candidate prey protein). *C)* Immunofluorescence analysis of the subcellular localization of the wild-type and SCO1 M294V bait proteins. Cell lines were fixed and stained with a mitochondrial marker HSPA9 (red), a nuclear marker DAPI (blue) and FLAG (green) to detect the SCO1 bait proteins. The yellow indicates mitochondrial co-localization. All images were acquired at 40X magnification. *D)* Dot plots depicting the distribution of both non-mitochondrial (grey) and mitochondrial (green) prey proteins as a function of the SAINT score. *E)* Dot plots showing the presence (green) or absence (red) of an interaction between SCO1 or SCO2 bait proteins and the indicated prey proteins.

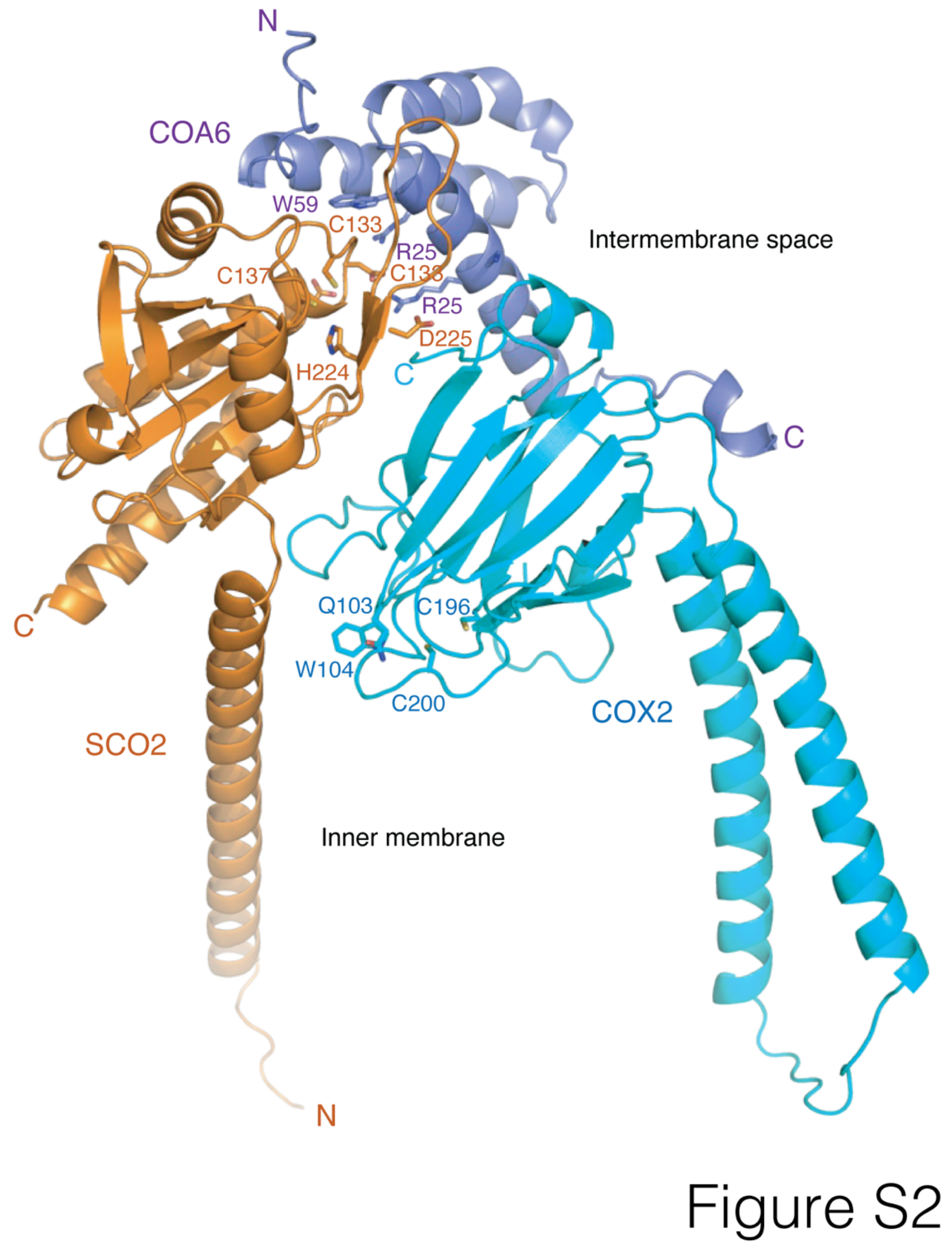


**Figure S2. Predicted model of the SCO2:COA6:COX2 complex.** Ribbon representation of SCO2 (orange), COA6 (lavender) and COX2 (light blue) in a similar orientation as the SCO1 and COA6 subunits in the SCO1:COX2:COX16:COA6:COX20 ternary complex (see Fig. 5C, for reference). The transmembrane domains of SCO2 and COX2 are at the bottom of the Figure. Side chains of SCO2 and

COA6 that are predicted to interact are shown as stick models and labelled appropriately. SCO2 Cu-ligand residues are also shown. Side chains of COX2 that make up the SCO2:COX2 interface and the $Cu_A$ site Cys residues are shown for comparison and labelled appropriately.

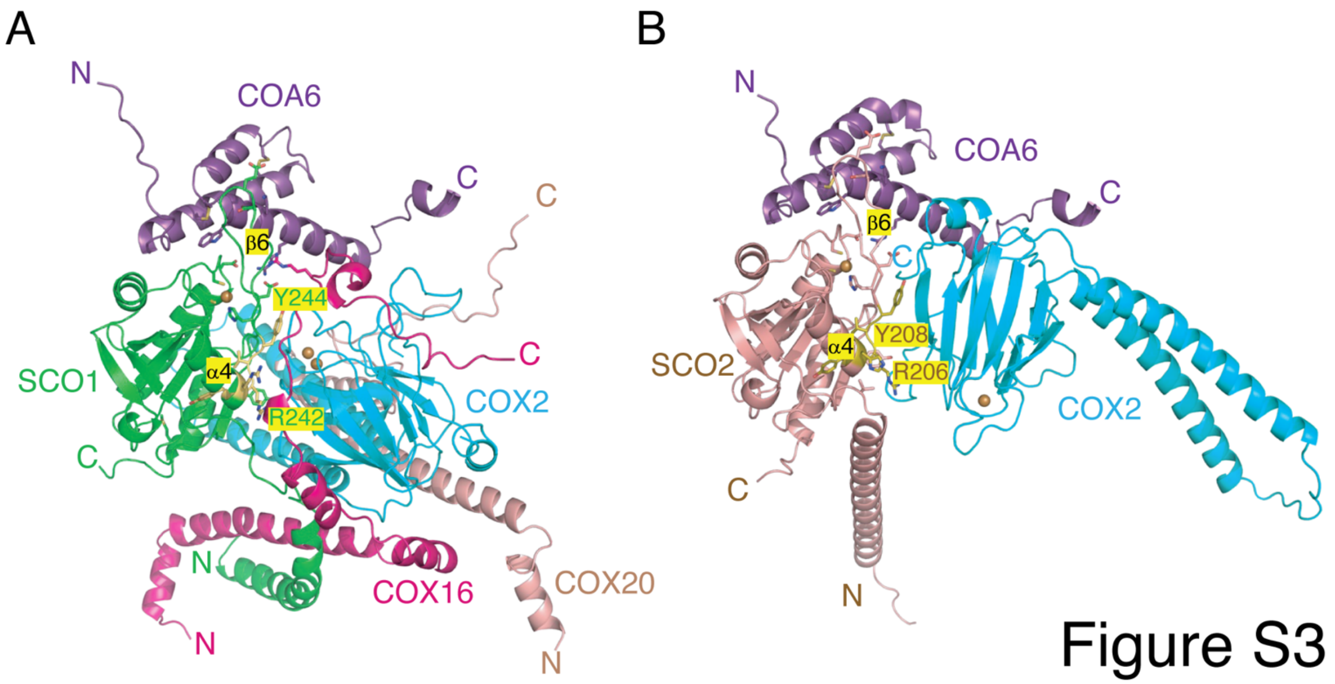


**Figure S3. Comparison of the predicted SCO1:COX16:COA6:COX2:COX20 and SCO2:COA6:COX2 structures.** *A)* The SCO1:COX16:COA6:COX2:COX20 complex is drawn as a ribbon diagram and coloured as in Fig. 4G. *B)* The SCO2:COA6:COX2 complex is drawn and coloured as in Fig. S2. For panels *A&B)***,** the complexes are shown in an identical orientation of SCO1/SCO2 and COA6. The three central conserved side chains of residues within a motif critical for yeast Sco1p (KKY**RVY**F) to interact with yeast Cox2p (4) are shown for both human SCO1 $^{239}$(RAY**RVY**Y)$^{245}$ and SCO2 $^{203}$(HSY**RVY**Y)$^{209}$ where they are predicted to similarly interact with COX2. Secondary structure elements on SCO1 and SCO2 before and after this motif are labeled appropriately (α4 and β6).

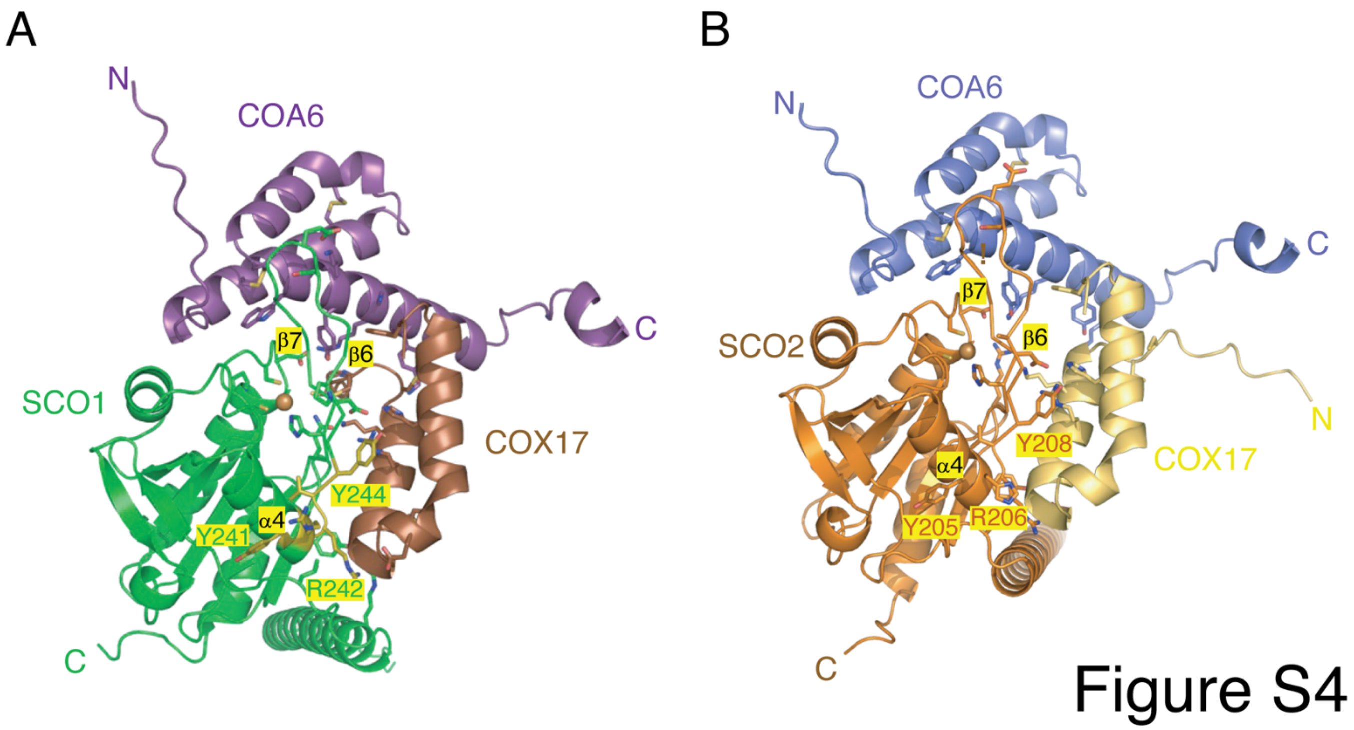


**Figure S4. Predicted SCO1:COA6:COX17 and SCO2:COA6:COX17 structures highlight a conserved interface critical for SCO interactions with COX17.** *A)* The SCO1 (green), COA6 (purple) and COX17 (brown) complex is drawn as a ribbon diagram in approximately the same orientation of SCO1 as depicted in Fig. 5C. *B)* The SCO2 (orange), COA6 (lavender) and COX17 (gold) complex is drawn in the same orientation as panel *A)*. For panels *A&B)***,** the complexes are shown in an identical orientation with respect to the interaction of each SCO with COX17. The three central conserved side chains of residues within a motif critical for yeast Sco1p (KKY**RVY**F) to interact with yeast Cox2p (4) are shown for both human SCO1 $^{239}$(RAY**RVY**Y)$^{245}$ and SCO2 $^{203}$(HSY**RVY**Y)$^{209}$ where they are predicted to similarly interact with COX17. Secondary structure elements on SCO1 and SCO2 before and after this motif are labeled appropriately (α4 and β6).

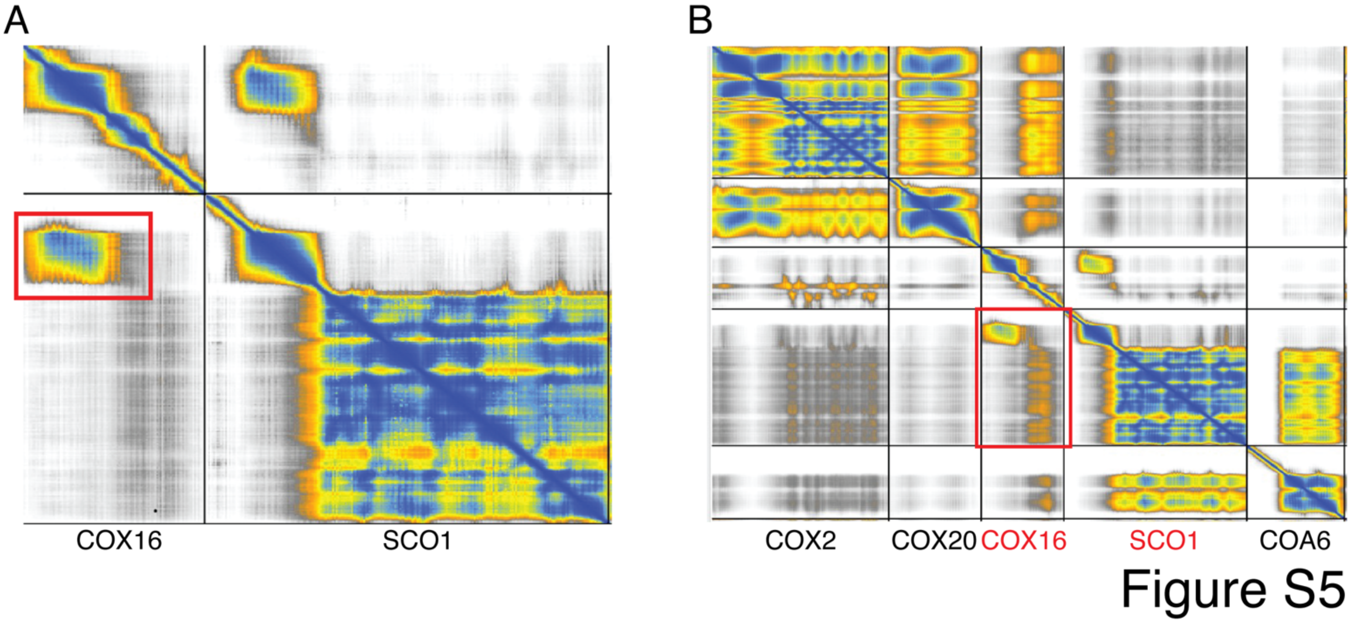


**Figure S5. The SCO1:COX16 interaction is stabilized primarily by their transmembrane domains.** Predicted Alignment Error matrices from AlphaFold3 for the *A)* binary and *B)* ternary complexes containing SCO1 and COX16. The PAE matrices are drawn with ChimeraX (5, 6).

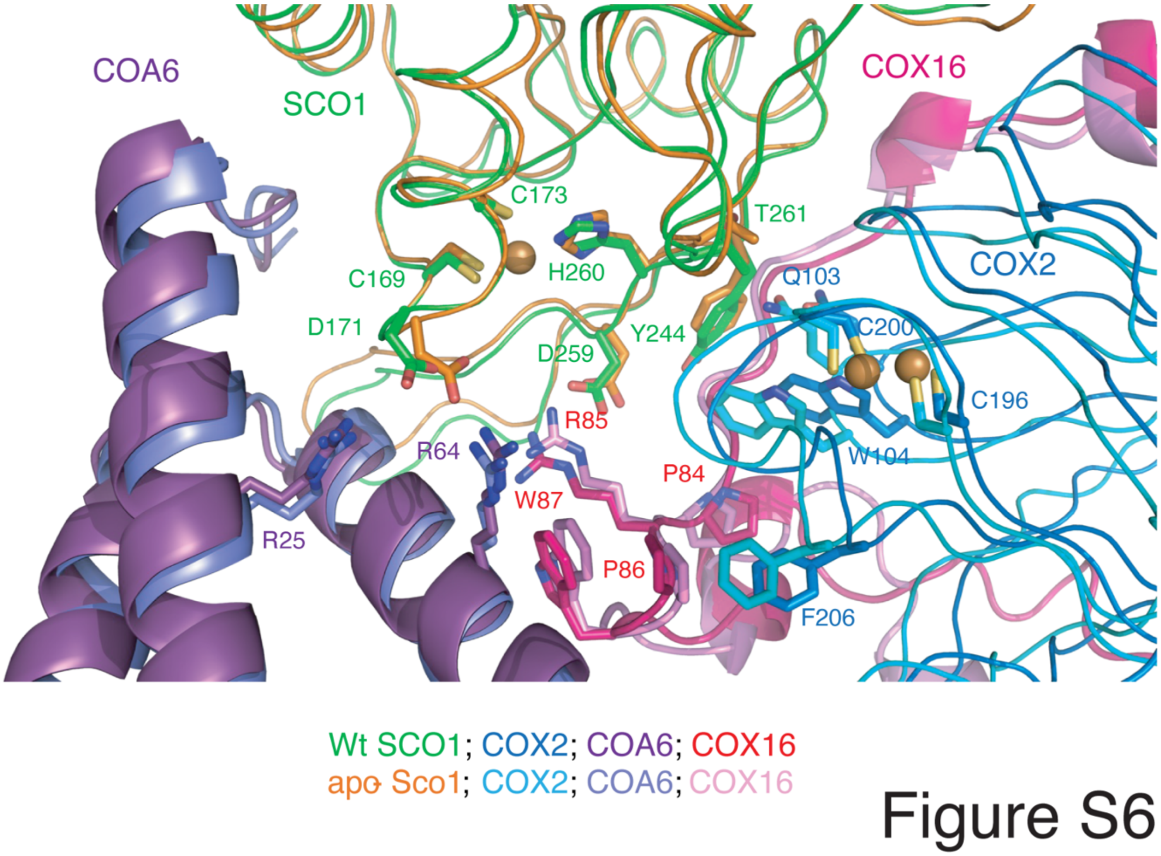


**Figure S6. Impact of SCO1 metalation state on the structure of the SCO1:COX16:COA6:COX2:COX20 ternary complex in the context of $Cu_A$ site maturation.**

View of wild-type $Cu^+$-loaded human SCO1 (holo-SCO1; green) in complex with COX16 (dark red), $Cu^+$-loaded COX2 (dark blue), and COA6 (dark purple) overlaid on the complex of wild-type apo-SCO1 (tawny), COX16 (pink-red), apo-COX2 (light blue) and COA6 (lavender). Drawn in a similar orientation as Fig. 5C, D and F. COX20 is not shown for the sake of clarity.

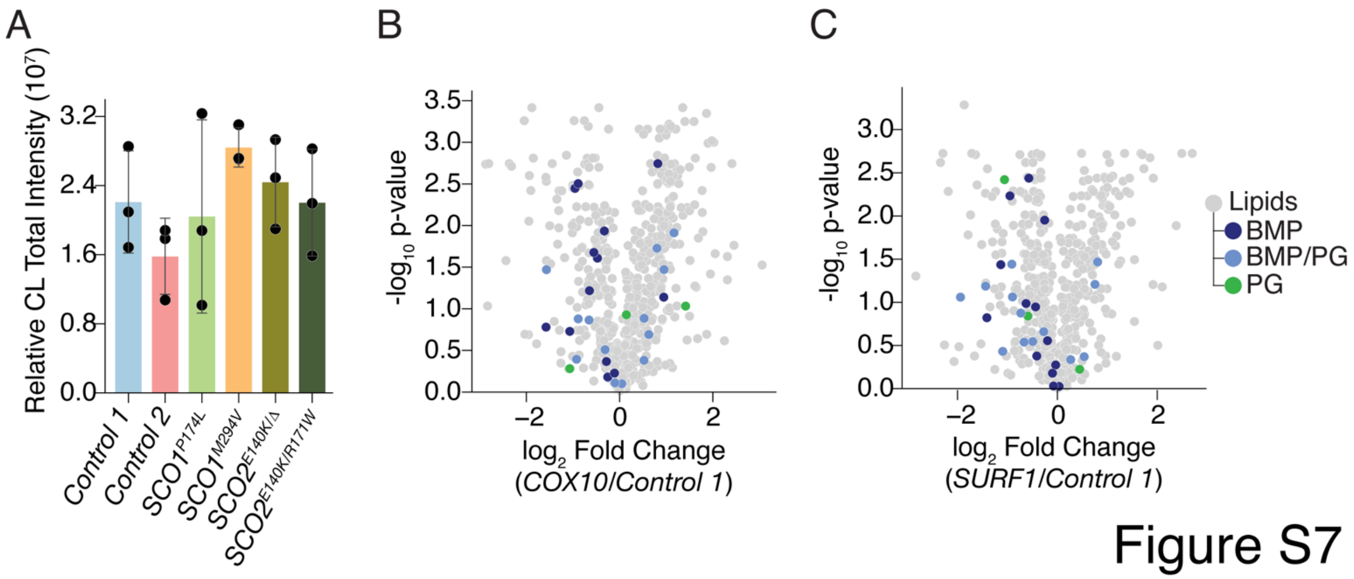


**Figure S7. BMP and PG lipids are not perturbed in other COX deficient patient backgrounds.** *A)* Combined relative intensities of all detected CL species in *SCO1* patient (P174L and M294V), *SCO2* patient (E140K), and *Control* fibroblast lines. *B)* Relative lipid abundance in *COX10* patient fibroblasts compared to *Control 1* fibroblasts, versus statistical significance. *C)* Relative lipid abundance in *SURF1* patient fibroblasts compared to *Control 1* fibroblasts, versus statistical significance. For panels *B&C)* Identified BMP and PG species are coloured in dark blue and green, respectively. Lipids that could only be identified as being either BMP or PG (BMP/PG) are coloured in light blue. For all panels, n=3 for each condition.
